# Cognitive outcome and its neural correlates after cardiorespiratory arrest in childhood

**DOI:** 10.1101/2023.05.02.539098

**Authors:** Sharon Geva, Aparna Hoskote, Maneet Saini, Christopher A. Clark, Tina Banks, Kling W. K. Chong, Torsten Baldeweg, Michelle de Haan, Faraneh Vargha-Khadem

## Abstract

Hypoxia-ischaemia (HI) can result in structural brain abnormalities, which in turn can lead to behavioural deficits in various cognitive and motor domains, in both adult and paediatric populations. Cardiorespiratory arrest (CA) is a major cause of hypoxia-ischaemia in adults, but it is relatively rare in infants and children. While the effects of adult CA on brain and cognition have been widely studied, to date, there are no studies examining the neurodevelopmental outcome of children who suffered CA early in life.

Here, we studied the long-term outcome of 28 children who suffered CA during infancy or childhood (i.e., before age 16). They were compared to a group of control participants (n = 28) matched for age, gender and socio-economic status. The patient group had impairments in the domains of memory, language and academic attainment (measured using standardised tests; impairment defined as a score > 1.5 standard deviations below the control group mean). Individual scores within the impaired range were most commonly found within the memory domain (79%), followed by attainment (50%), and language (36%). The patient group also had reduced whole brain grey matter volume, and reduced volume and fractional anisotropy of the white matter.

In addition, lower performance on memory tests was correlated with bilaterally reduced volume of the hippocampi, thalami, and striatum, while lower attainment scores were correlated with bilateral reduction of fractional anisotropy in the superior cerebellar peduncle, the main output tract of the cerebellum.

We conclude that patients who suffered early CA are at risk of developing specific cognitive deficits associated with structural brain abnormalities.

## 1 Introduction

Neonatal hypoxia-ischaemia (HI) can have a circulatory or respiratory origin (Fatemi et al., 2009; Thompson et al., 2014b; van Schie et al., 2007; Young et al., 2015), and its adverse effects on brain structure and function are mediated through various molecular pathways (Fatemi et al., 2009). It is as yet unknown whether HI of different aetiological origins can result in different outcomes. However, the most common patterns of brain injury associated with HI in term neonates include damage to: (i) the hippocampus which underpins cognitive memory and learning, the basal ganglia (specifically the neostriatum; comprised of the caudate nucleus and the putamen) which support habit formation and skill learning, and the thalamus (Cowan et al., 2003); and, (ii) the watershed zones, which usually affects the white matter, although, in more severely affected children, can also cause damage to the surrounding cortex (Fatemi et al., 2009).

These brain structures affected by early HI form part of various functional and structural neural circuits, and therefore, damage to these structures can result in behavioural sequelae in various developmental domains, including motor and cognitive functions. The developmental outcome can vary in severity, from an overt neurological outcome, such as cerebral palsy, through subtle cognitive abnormalities, to normal development (Fatemi et al., 2009). Adverse cognitive outcomes, especially affecting memory function, have been reported by our group after diverse aetiologies of neonatal hypoxia-ischaemia (Gadian et al., 2000; Vargha-Khadem et al., 1997). In two subsequent studies, we examined the association between specific HI aetiology and memory impairments. The two cohorts included children treated for acute hypoxemic respiratory failure (AHRF; Cooper et al., 2015), and children treated at birth for transposition of the great arteries (TGA; Muñoz-López et al., 2017) using a corrective arterial switch operation. A substantial number of patients (45% in Cooper et al., 2015; 33% in Muñoz-López et al., 2017) showed marked memory impairments, and on the group level, this was associated with significant reduction in bilateral hippocampal volume, on a background of otherwise normal cognitive and neurological outcome.

Cardiorespiratory arrest (CA) is another well-known cause of HI. The cognitive and motor function following CA in adults has been studied before (see below). However, CA is relatively rare in children as compared to adults (Engdahl et al., 2003; Holmberg et al., 2019; Tsao et al., 2022), and this may partly explain the reason for the scarcity of data on functional outcomes following CA in children. In addition, in children, cardiac arrest usually has different aetiology than in adults: the most common cause of paediatric cardiac arrest is hypoxia and rarely a primary cardiac event, unlike in adults where the common cause is ventricular fibrillation (Manole et al., 2009; Tress et al., 2010). Moreover, during infancy and childhood neuronal maturation and synaptogenesis still occur. As a consequence, the sequelae of childhood cardiac arrest and the recovery patterns could differ from those seen in adults (Tress et al., 2010).

We, therefore, studied a rare cohort of children and adolescents who had suffered from CA, with the aim of understanding their long-term cognitive outcome in relation to the integrity of known neuroanatomical targets of HI.

In adults, it is now well-established that some patients who had cardiac arrest suffer mild (33-66% of survivors) or severe (7-28%) memory impairments (Alexander et al., 2011; Bunch et al., 2004; Cronberg et al., 2009; Grubb et al., 1996; Mateen et al., 2011; Moulaert et al., 2009; O’Reilly et al., 2003; Sauve, 1995; Sunnerhagen et al., 1996; Tiainen et al., 2007); (but see Bertini et al., 1990). A review concluded that performance in all cognitive domains can be compromised (Moulaert et al., 2009), with executive functions most commonly affected (Cronberg et al., 2009; Drysdale et al., 2000; Lim et al., 2004; Nunes et al., 2003). Lim et al. (2004) found no patients with isolated memory impairment, arguing that following CA, memory impairments always co-occur with other cognitive dysfunctions. When they occur, the memory impairments are chronic and affect recognition as well as recall, suggesting non-focal brain injury (Drysdale et al., 2000; Grubb et al., 2000). However, verbal long term memory was found to be more affected than visual long term memory (Grubb et al., 1996). It is well-documented that impairments of memory and other cognitive functions affect independence, and quality of life of both patients and carers (Pusswald et al., 2000).

In paediatric populations, two large cohort studies examined general cognitive outcome 12 months after CA, using the Mullen Scales of Early Learning for children below 6 years of age, or two subscales from the Wechsler Abbreviated Scale of Intelligence (WASI) for older children. Most patients who had out-of-hospital CA (total n = 60) performed within the impaired range (Slomine et al., 2016), while among those who had in-hospital CA (total n = 100), less than half showed similar impairment (Slomine et al., 2018a). Further testing of the older children (> 6 years of age; n = 41), revealed that some of the patients had impairments in verbal recall (∼ 20%, tested with the California Verbal Learning Test), visual recall (∼ 50%, Rey–Osterrieth complex figure), verbal fluency (∼ 40%), and motor functioning (∼ 40%, Grooved Pegboard and Beery-Buktenica Visual-Motor Integration) (Slomine et al., 2018b). A single small study of cardiac arrest in adolescents found that 6 out of 8 patients whose memory status could be assessed showed mnemonic impairments early after the event. In a follow-up examination (minimum 6 months post CA) two of these adolescents fully recovered, and the remaining four had residual deficits (Maryniak et al., 2008).

Medical factors that influence long-term outcome following cardiac arrest in adults include being awake on admission and a short duration of coma, both predict better long-term cognitive functioning (Moulaert et al., 2009), whereas duration of cardiac arrest has been reported to be a reliable predictor of memory impairment (Grubb et al., 1996). Other medical and demographic variables were not associated with cognitive outcome (e.g., time of arrival of ambulance (Moulaert et al., 2009; Sunnerhagen et al., 1996); call-to-shock time (Mateen et al., 2011), whether the arrest was witnessed or not (Mateen et al., 2011); bystander cardiopulmonary resuscitation (Mateen et al., 2011); location at time of cardiac arrest (O’Reilly et al., 2003); resuscitation variables and co-morbidities (Moulaert et al., 2009); sex, age at cardiac arrest and time since event (Mateen et al., 2011; Moulaert et al., 2009); or therapeutic hypothermia (Tiainen et al., 2007)).

Studies of paediatric population yielded similar results, showing that therapeutic hypothermia was not associated with functional outcome 12 months post CA, regardless of CA location (Moler et al., 2017, 2015; Scholefield et al., 2018). However, factors related to the CA and resuscitation were associated with neurobehavioural outcome (Ichord et al., 2018; Meert et al., 2018, 2016). For example, worse functional outcome was predicted by having out-of-hospital arrest (versus in-hospital), arrest of respiratory origin (versus cardiac origin), and longer duration of chest compressions (Ichord et al., 2018). Older age at CA was also associated with poorer functional outcome (Slomine et al., 2018a, 2016).

We therefore studied the long-term cognitive outcome of a cohort of children and adolescents who suffered a cardiorespiratory arrest early in life. Based on the behavioural results from adult studies, and from studies of children who suffered hypoxia-ischaemia of other aetiologies (Cooper et al., 2015; Muñoz-López et al., 2017), we hypothesised that impairments would be found primarily in the domain of cognitive memory. Furthermore, following findings that early language development and general cognitive outcome (Slomine et al., 2018a, 2016), as well as later verbal fluency and recall (Slomine et al., 2018b) are affected by CA, we also examined detailed language abilities and general academic attainment. We hypothesised that age at CA, duration of CA and receiving ECMO treatment would be reliable predictors of cognitive outcome. Lastly, we examined whether the patient cohort had grey matter injury to structures commonly affected by HI (i.e., the dorsal striatum of the basal ganglia which includes the caudate nucleus and putamen, the hippocampus and the thalamus), and whether this damage is associated with specific domains of cognitive dysfunction. We also studied the status of global white matter, as well as the integrity of white matter tracts associated with the cognitive functions measured here, including the fornix in relation to memory (Dogan et al., 2022), the arcuate fasciculus/superior longitudinal fasciculus (AF/SLF) and inferior longitudinal fasciculus/Inferior fronto-occipital fascicle (ILF/IFOF) in relation to language (Dick et al., 2014; Friederici, 2015), and the superior cerebellar peduncle in relation to attainment (Albazron et al., 2019).

Based on previous studies of paediatric populations, we hypothesised that: (i) specific aspects of memory function will be associated with damage to the hippocampus and fornix (Cooper et al., 2015; Mishkin et al., 1998; Muñoz-López et al., 2017; Vargha-Khadem et al., 1997), and the thalamus (Sweeney-Reed et al., 2021); (ii) language function will be associated with damage to the basal ganglia (Martinez-Biarge et al., 2012b, 2012a, 2010), and/or the major white matter language tracts (AF/SLF and ILF/IFOF; Friederici, 2015); and, (iii) general attainment outcome will be associated with diffusivity parameters of the superior cerebellar peduncle, the major efferent pathway of the cerebellum which sends information to the cortex through the thalamus (Albazron et al., 2019).

## 2 Materials and methods

The study was approved by the London-Bentham Research Ethics Committee (REC Reference No. 05/Q0502/88) and all participants and their carers read an age-appropriate information sheet and gave written informed consent according to the Declaration of Helsinki, before commencing the study.

### 2.1 Participants

From a database of patients who suffered a substantial cardiorespiratory arrest (CA) and were seen in Great Ormond Street Hospital between the years 1997 and 2014, we initially included all those who would have been 8-22 years of age at the time of study participation. Inclusion criteria were: no history of other developmental or genetic disorders or conditions which may affect neurological function, living in the UK, both patient and parent/guardian are English-speaking, and MRI compatibility. Out of 326 patients, 182 (55%) were deceased, and a further 91 were excluded based on the above criteria. Among the 53 patients who were included, 25 did not participate in the study (13 refused; 11 had missing contact details; 1 agreed but did not attend). The remaining 28 patients participated in the study. For full recruitment details see Supplementary Figure 1. For clinical and demographic details of each participating patient see Supplementary Table 1. The following medical variables describe the patient cohort: age at CA (mean: 3.4 ± 4.4 years), total arrest time (mean 10.8 ± 19.8 min; coded as 1 min in 3 cases where medical notes stated ‘short downtime’), Extracorporeal membrane oxygenation (ECMO) treatment (14 yes / 14 no), number of days on ECMO (n = 14; mean: 9 ± 4.11), CA location (1 out of hospital – at home / 27 in hospital), mechanical circulatory support with ventricular assist device (1 yes / 27 no), and heart transplant (6 yes / 22 no).

To evaluate the representativeness of the recruited sample, we compared demographic and clinical variables of the patients who participated in the study, to those of the 25 patients who met the inclusion criteria but did not participate. The two patient groups did not differ on any of the variables (see Supplementary Table 2), suggesting that assessment of only half of the eligible patients did not bias our final sample with regard to medical and demographic variables.

Thirty healthy participants were recruited mainly from schools in the London area to act as a comparison control group. Inclusion criteria were age 8-22 years at the time of study participation, no relevant medical history, no developmental disorders, term birth, and both child and parent are English-speaking. Inclusion was determined based on answers given by parents to a detailed medical questionnaire administered via telephone. Two healthy controls were later excluded due to abnormal findings on the MRI scan, resulting in a control group of 28 participants.

Socio-Economic-Status (SES) of participants was defined using the National Office of Statistics data (Income Deprivation Affecting Children Index; IDACI). Data were missing for two patients who live outside of England, and for one healthy participant whose post code was not obtained. Both groups were representative of the general population (one sample t-test, p > 0.05 for both groups). The group of patients did not differ from the control group in age, gender, handedness (Oldfield, 1971), or SES (see Table 1). Group demographic and clinical information is provided in Table 1 and Supplementary Table 2.

**Table 1.** Demographic information of the groups of patients and healthy controls.

| Variables |  | Controls | Patients | Statistical test | p-value |
| --- | --- | --- | --- | --- | --- |
| **Age at first CA (days)** | Mean (SD) | N/A | 1242 (1614) | N/A | N/A |
|  | Range |  | 0 - 5031 |  |  |
| **Time between CA and test (years)** | Mean (SD) | N/A | 8.5 (3.9) | N/A | N/A |
|  | Range |  | 1 - 15 |  |  |
| **Age at test** | Mean (SD) | 12.4 (3.2) | 12.4 (3.3) | t-test | 0.973 |
|  | Range | 8 - 21 | 8 - 20 |  |  |
| **Gender** | F | 18 | 17 | Chi square | 0.783 |
|  | M | 10 | 11 |  |  |
| **SES IDACI rank (k)** | Mean (SD) | 16 (10.5) | 17 (9.4) | t-test | 0.833 |
|  | Range | 1 - 32.3 | 0.6 - 30.3 |  |  |
| **SES IDACI decile** | Mean (SD) | 5.6 (3.13) | 5.8 (2.92) | Mann-Whitney | 0.795 |
|  | Range | 1 - 10 | 1 - 10 |  |  |
| **Handedness** | Right | 25 | 25 | Chi square | 0.717 |
|  | Left | 1 | 2 |  |  |
|  | Ambidextrous | 2 | 1 |  |  |

CA = cardiac arrest; IDACI = Income Deprivation Affecting Children Index; SES = Socio-Economic-Status

### 2.2 Neuropsychological assessment

All participants completed a battery of standardised cognitive tests at the UCL Great Ormond Street Institute of Child Health. For most participants assessments were completed over one day. Neuropsychological assessments took around 3.5 hours in total, usually with a lunch break in the middle. Participants were given as many breaks as needed. The following cognitive tests were administered: (a) **Wechsler Abbreviated Scale of Intelligence** (WASI; Wechsler, 1999): measuring verbal IQ (VIQ; based on 2 subtests) and performance IQ (PIQ; based on 2 subtests), and generating a full-scale IQ index score (FSIQ; based on the 4 subtests); (b) **The Children Memory Scale** (CMS; ages 8-16; Cohen, 1997) or the **Wechsler Memory Scale** (WMS; ages 16 and above; Wechsler, 2009): measuring immediate and delayed verbal and visual memory, general memory and recognition memory; (c) **Rivermead Behavioural Memory Test** (RBMT; children version for ages 8-10; version II for ages 11 and above; Wilson et al., 1991): measuring everyday episodic memory; (d) **Wechsler Individual Achievement Test – II** (WIAT-II; Smith, 2001): measuring academic attainments. Four subtests were used from this test battery: Numerical Operations, Word Reading, Reading Comprehension, and Spelling. (e) The **Expression, Reception and Recall of Narrative Instrument** (ERRNI; Bishop, 2004): measuring language ability using a narrative test, and generating five scores: (i) Ideas 1: amount of information given in initial storytelling (while looking at a picture book); (ii) Ideas 2: amount of information given in the second storytelling (after 30 min delay, without looking at the picture book); (iii) Forgetting level (between the two storytelling sessions); (iv) Mean Length of Utterance (MLU; averaged across both storytelling sessions); (v) Comprehension score: accuracy of answers given to a number of questions presented to the participant while looking at the pictures, at the end of the testing session. (f) The **Sunderland Memory Questionnaire for Children and Adolescents** (Sunderland et al., 1983) was completed by the participants’ primary carer.

### 2.3 Imaging data acquisition

MR images were acquired using a 1.5T Siemens Avanto (Germany) MRI scanner at Great Ormond Street Hospital, London. For most participants the MRI scan was performed on the same day as the neuropsychological assessments. The following scans were acquired: T1-weighted Fast Low Angle Shot (FLASH) scan (repetition time (TR) = 11 ms, echo time (TE) = 4.94 ms, flip angle = 15°, field of view = 224 × 256 mm, 176 slices, sagittal plane, voxel size: 1 x 1 x 1 mm); T2-weighted scan (TR = 4920 ms, TE = 101 ms, flip angle = 90°, field of view = 220 × 256 mm, 25 slices, voxel size: 7 x 6 x 4 mm); and a Diffusion Weighted Image (DWI; TR = 7300 ms, TE = 81 ms, field of view: 240 x 240 mm, 60 slices, axial plane, voxel size: 2.5 x 2.5 x 2.5 mm, 60 gradient directions, b-value: 1000 s/mm²). Four patients did not have an MRI scan (one found to be non-MRI compatible prior to scanning, one stopped the scan in the middle, for one the parent declined the scan following testing, and for one patient the MRI data were unusable because of artefacts created by braces).

One of the authors, a consultant paediatric neuroradiologist (WKKC) visually inspected all T1-weighted and T2-weighted MRI scans, while being blinded to participants’ diagnosis or group affiliation. The following pre-defined brain structures were rated as either normal or small on inspection: hippocampus, mesial-temporal lobe, fornix, dorsomedial thalamus, mammillary bodies, basal ganglia, and cerebellum. The size of the mammillary bodies was evaluated in relation to two internal landmarks: the optic chiasm and the anterior commissure. The lateral ventricle was rated for being normal or enlarged/dilated. Periventricular white matter was rated for amount (normal or reduced), and for presence of abnormalities such as scars. The size of the genu and the splenium of the corpus callosum was compared, where splenium > genu was rated as normal; while a reversed relationship (genu > splenium) was rated as abnormal. Other abnormalities such as ischaemic or atrophic damage, haematoma or gliotic scars were noted in 11 patients. The scans of all healthy controls were rated as normal. See Supplementary Table 3 for radiological findings of all the patient participants.

### 2.4 Imaging data pre-processing

#### 2.4.1 Processing of T1-weighted images

T1-weighted FLASH images were processed using the Statistical Parametric Mapping software (SPM12; Wellcome Centre for Human Neuroimaging, London, UK; https://www.fil.ion.ucl.ac.uk/spm/) running in the Matlab environment (2021a Mathworks, Sherbon, MA, USA). Images were segmented into grey matter (GM), white matter (WM), cerebro-spinal fluid (CSF) and other tissue type based on probability maps, using the Segment tool. This procedure combines tissue segmentation, bias correction, and spatial normalization in a single unified model (Ashburner and Friston, 2005). Normalisation was carried out using Diffeomorphic Anatomical Registration Through Exponentiated Lie Algebra (DARTEL, Ashburner, 2007), which creates templates and normalises individual scans to MNI space based on all participants’ images (n = 52). Images were smoothed using an 8 mm Full-Width at Half Maximum (FWHM) Gaussian kernel. Tissue volumes were calculated using the Tissue Volume utility in SPM12, and intra-cranial volume (ICV) was defined as a combination of whole brain GM, WM and CSF volumes.

#### 2.4.2 Processing of Diffusion Weighted Images

Processing of DWI was carried out using the FMRIB’s Software Library (FSL; http://www.fmrib.ox.ac.uk/fsl, Smith et al., 2004). First, eddy current distortions and head movements were corrected by registering all DTI volumes to the first b0 volume, using the eddy correct tool. Next, the FSL Brain Extraction Tool (BET) was applied to obtain a binary brain mask for each participant. A diffusion tensor model was fitted at each voxel using DTIFIT to obtain the eigenvalues (λ1, λ2, λ3) and eigenvectors (V1, V2, V3) of the diffusion tensor matrix. Maps of Fractional Anisotropy (FA), Mean Diffusivity [MD = (λ1+λ2+λ3)/3], and Radial Diffusivity [RD = (λ2+λ3)/2] were generated based on these eigenvalues. Each dataset was visually inspected for data quality, and those with severe artefact were discarded from further analysis. The data from one patient and one healthy volunteer were discarded because of motion artefacts created by the participant’s own movement. Four additional datasets (two patients and two healthy volunteers) were discarded due to motion artefacts related to table motion during data acquisition. We then applied nonlinear registration of each image into standard space provided by FSL. Lastly, a mean image was created based on data from the remaining participants (n = 46) and skeletonised for the whole group. It was thresholded with the default threshold value of 0.2. Masks were created for studying specific tracts of interest using the JHU Atlas (Mori et al., 2005), binarized, and mean FA, MD and RD values were extracted from the tracts of each participant.

#### 2.4.3 Volumetry of subcortical structures

We performed automated segmentation of the subcortical structures (bilateral hippocampus, thalamus, caudate nucleus and putamen as regions of interest, brain stem as control region) using FSL-FIRST v.6.0 (Patenaude et al., 2011) from all participants’ T1-weighted MRIs in native space. Volumes were segmented in each hemisphere and visually inspected for accuracy. One control participant was excluded for poor registration and outlier values for ICV and all subcortical structures (outliers defined as values > Mean + 3SD of the control group). Volumetric measurements were normally distributed (Shapiro-Wilk test, p > 0.05 for all).

### 2.5 Data Analysis

#### 2.5.1 Behavioural data analysis

For dimensionality reduction, we ran principal component analyses (PCAs) on the behavioural scores of the different tests (following Corbetta et al., 2015; Ramsey et al., 2017; Salvalaggio et al., 2020) within three domains: (i) Memory (Sunderland, CMS/WMS verbal delayed score, RBMT and ERRNI Forgetting); CMS/WMS visual delayed scores were not included as a previous study by Buck et al. (2021) suggested that as memory load in this test is low, the score is not sensitive enough to small variability in delayed memory abilities; and in line with an adult study, showing that verbal long term memory was more affected than visual long term memory (e.g. Grubb et al., 1996); (ii) Language (ERRNI scores other than Forgetting); and, (iii) Attainment (WIAT subtests). In order to maximize the use of the data, we eliminated the WIAT reading comprehension scores from the final PCA, as 12 participants did not complete this subtest. However, the components derived from four versus three WIAT subtests were highly positively correlated (Pearson’s r = 0.98, p < 0.001). IQ scores highly correlated with all other scores and were therefore excluded from the PCAs.

We used oblique rotation, because we assumed the components are not orthogonal. Components had to satisfy two criteria: (i) the eigenvalues had to be > 1; (ii) the percentage of variance accounted for had to be > 10% (Corbetta et al., 2015; Ramsey et al., 2017; Salvalaggio et al., 2020). For each domain, we obtained one component that accounted for the majority of the variance. Although our sample size was relatively small for PCA analysis, all criteria indicated adequacy of the method: Kaiser–Meyer–Olkin measure of adequacy was above 0.5, Bartlett’s test of sphericity was significant, and extraction values were all equal to, or higher than 0.3. See Supplementary Table 4 for further details on the three PCAs. We then compared the patient and control groups on behavioural scores using independent sample 1-tailed t-tests.

Within the patient group, we examined whether behavioural outcome was associated with cardiac arrest factors (age at CA and total arrest time; 1-tailed Spearman’s rho, threshold p = 0.008 following Bonferroni’s correction for multiple comparisons), or ECMO treatment (Mann-Whitney U Test).

#### 2.5.2 Group differences in tissue volumes and microstructure

To examine group differences (patients versus controls) in tissue volumes and microstructure we performed a series of MANOVAs, and we report main effects of group and interaction effects. We then established that effects remained the same when controlling for age at test, by running MANCOVAs, with age at test as a covariate. MANOVAs were followed by post-hoc independent sample 1-tailed t-tests.

MANOVAS were performed for the following (i) 2x3 MANOVA of whole-brain tissue volume: GM, WM and CSF; (ii) 2x3 MANOVA of whole-brain diffusion parameters: FA, MD and RD; (iii) 2x4x2 MANOVA of subcortical grey matter structures of interest (four subcortical structures: hippocampus, thalamus, caudate nucleus, putamen; and two hemispheres: left and right); (iv) 2x3x2 MANOVA of white matter tracts of interest (three tracts: AF/SLF, ILF/IFOF, superior cerebellar peduncle (SCP); and two hemispheres: left and right). We also compared the two groups for FA values of the fornix (which was not included in the MANOVA as TBSS does not distinguish right and left hemispheric portions of the fornix).

We then examined the association between global tissue parameters (GM, WM and CSF volume, and FA, RD and MD) and cardiac arrest factors (total arrest time and age at CA covaried for age at test; 1-tailed Spearman’s rho, threshold p = 0.004 following Bonferroni’s correction for multiple comparisons), or ECMO treatment (Mann-Whitney U Test).

#### 2.5.3 Structure-function relationships

We aimed to establish a relationship between medical variables, brain abnormalities and cognitive performance, using Pearson’s correlations, and Benjamini–Hochberg FDR corrections for multiple comparisons (Benjamini and Hochberg, 1995). We only report correlations which were significant also when controlling for age at testing using partial correlation.

The dependent variables were the components obtained in the three PCAs of the behavioural performance (i.e. memory, language and attainment). Independent variables included: (i) Duration of cardiac arrest (as studies of adult cardiac arrest associated this variable with cognitive outcome); (ii) White matter abnormality: extracted mean FA for each participant in the fornix, and left and right AF/SLF, ILF/IFOF and SCP (Matejko and Ansari, 2015; Moeller et al., 2015); and, (iii) Grey matter abnormality: bilateral volumes of the hippocampus, thalamus, caudate nucleus and putamen. All data analyses were carried out in IBM SPSS Statistics for Windows (version 27.0, IBM Corp., Armonk, New York, USA).

## 3 Results

### 3.1 Behavioural performance

Patients had significantly lower scores compared to the control group, on all three components (independent sample t-test, Memory: t_(29.01)_ = 6.11, p < 0.001, Cohen’s d = 0.75; Language: t_(41.9)_ = 3.03, p = 0.004, Cohen’s d = 0.94; Attainment: t_(49)_ = 4.38, p < 0.001, Cohen’s d = 0.86). See Figure 1. Age at cardiac arrest, its duration or ECMO treatment were not associated with behavioural scores (p > 0.05 for all). Supplementary Table 4 provides group statistics and number of participants who performed within the impaired range, in each of the subtests.

**Figure 1.**
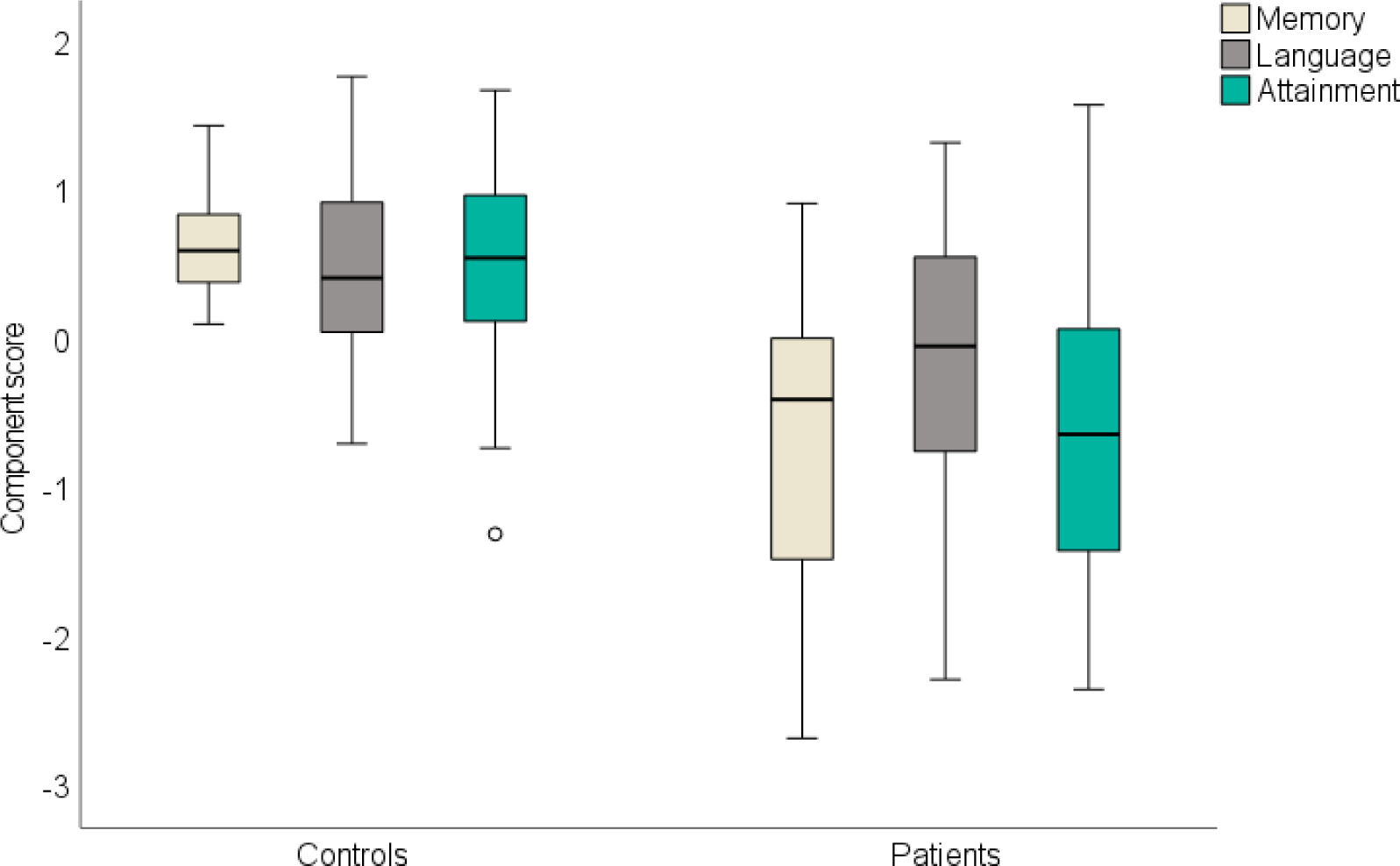
Group differences in cognitive outcome. Y axis represents the factor scores. The box represents the interquartile range (IQR), and the hinges represent the minimum and maximum values (excluding outliers). Small circles represent values which are > 1.5 times the IQR, below Q1 or above Q3.

### 3.2 Group differences in tissue volume and microstructure

#### 3.2.1 Whole brain measurements

Whole brain measurements in each group can be seen in Figure 2. A 2x3 MANOVA of whole brain tissue type volume revealed a significant main effect of group (F_(1,50)_ = 8.16, p = 0.006), and significant interaction between group and tissue type volume (F_(2,50)_ = 3.64, p = 0.043), effects which remained significant when controlling for age. Post-hoc tests showed that patients had lower GM and WM volumes (t_(50)_ = 3.14, p = 0.002; t_(50)_ = 2.61, p = 0.006; for GM and WM respectively), but there was no group difference in CSF volume (t_(50)_ = 0.59, p = 0.277).

**Figure 2.**
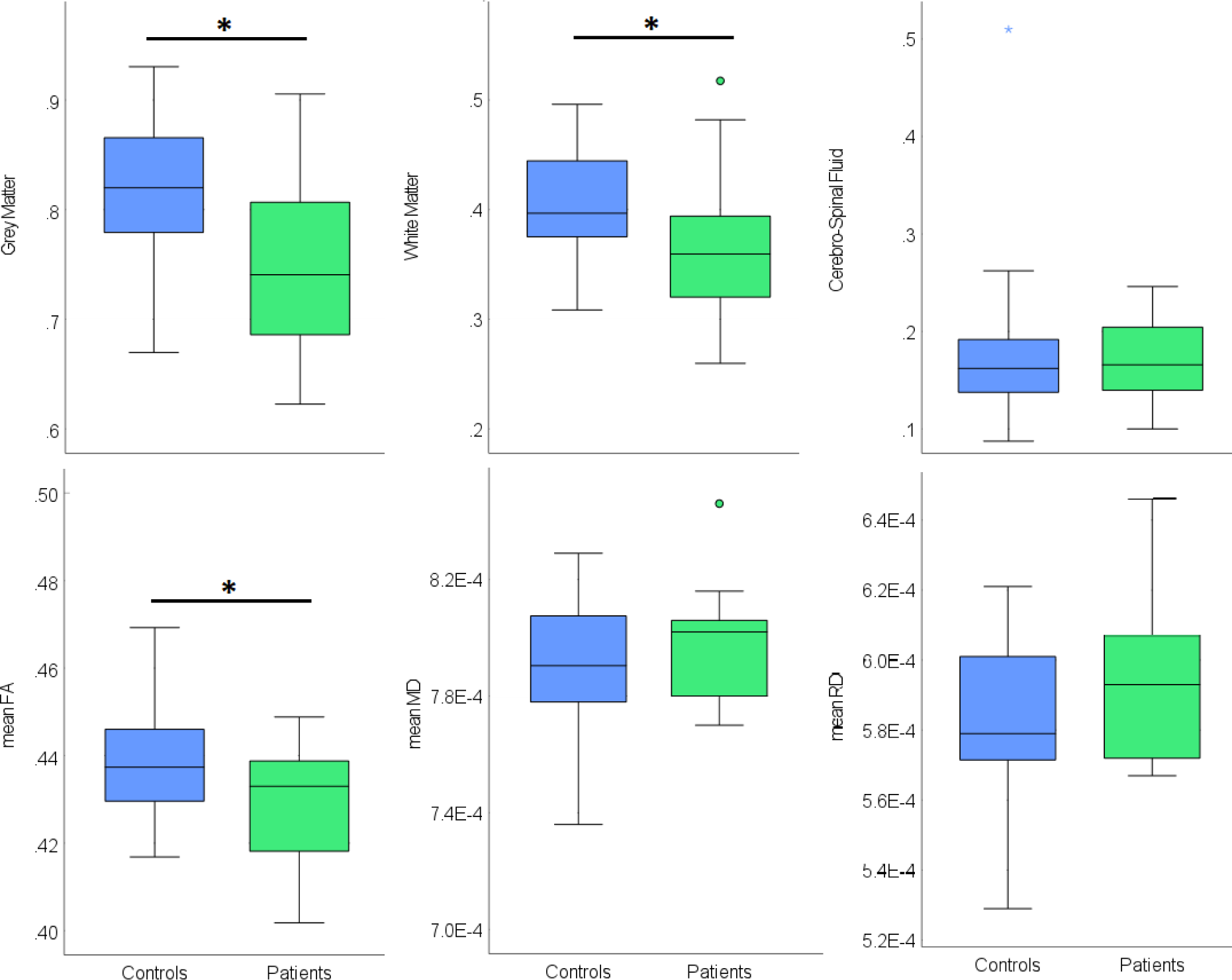
Group differences in whole brain measurements. Top: Grey matter, white matter and cerebrospinal fluid volumes (litre); Bottom panel: mean fractional anisotropy (FA), medial diffusivity (MD) and radial diffusivity (RD); in the control group (blue) and the patient group (green). Stars represent outliers, defined as values which are > 3 times the IQR either above Q3 or below Q1. Full circles represent values which are > 1.5 times the IQR. Black asterisk indicates significant group difference.

Similarly, a 2x3 MANOVA of whole brain diffusion parameters revealed a significant main effect of group (F_(1,43)_ = 5.24, p = 0.027), and significant interaction between group and diffusion parameter (F_(2,43)_ = 5.22, p = 0.027); once again, effects remained significant when controlling for age. Post-hoc tests showed that patients had significantly lower overall FA (t_(43)_ = 2.29, p = 0.014), but not MD or RD (t_(43)_ = 0.89, p = 0.189; t_(43)_ = 1.41, p = 0.084; for MD and RD respectively).

Older age at CA (covaried for age at test), longer arrest time and having ECMO treatment were not associated with whole brain parameters.

#### 3.2.2 Structures of interest

Having established that patients have lower whole brain white matter integrity (i.e., smaller volume and lower FA) and lower whole brain grey matter volume, we examined structures of interests. A 2x4x2 MANOVA of subcortical structures volumes (hippocampus, thalamus, caudate nucleus and putamen) revealed a significant main effect of group (F_(1,49)_ = 7.39, p = 0.009), and significant interactions between structure and group (F_(3,49)_ = 3.33, p = 0.021), and structure and hemispheres (F_(3,49)_ = 7.92, p < 0.001), but not between group and hemisphere (F_(1,49)_ = 0.27, p = 0.605). Post-hoc tests showed that patients had significantly bilateral reduction of the volume of the caudate nuclei and thalamus (t_(49)_ = 3.14, p = 0.002; t_(49)_ = 2.54, p = 0.007; for left and right caudate; t_(49)_ = 2.66, p = 0.006; t_(49)_ = 2.86, p = 0.003; for left and right thalamus), and significantly smaller left hippocampus (t_(50)_ = 3.21, p = 0.001), see Figure 3. The volume of the other subcortical structures did not significantly differ between patients and controls (p > 0.05 for all), nor did the volume of the brainstem, which acted as control structure (p > 0.05).

**Figure 3.**
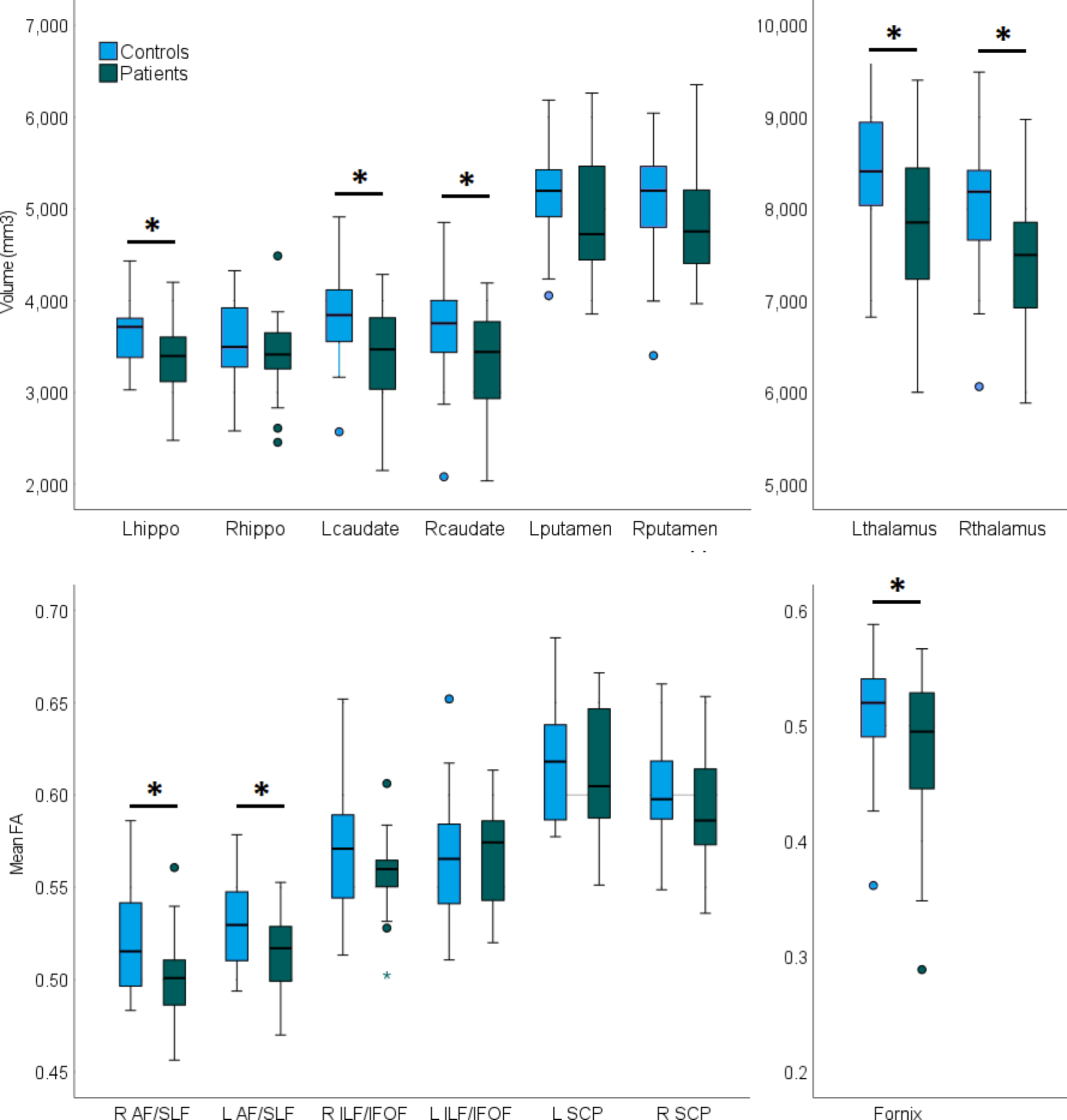
Group differences in structures of interest. Top: Volumes of the hippocampus (hippo), caudate nucleus (caudate), putamen and thalamus. Bottom: Mean FA of the AF/SLF, ILF/OFOF, SCP and fornix. Stars represent outliers, defined as values which are > 3 times the IQR either above Q3 or below Q1. Full circles represent values which are > 1.5 times the IQR. Black asterisk indicates significant group difference.

In a 2x3x2 MANOVA of FA in white matter tracts of interest (AF/SLF, ILF/IFOF and SCP), the effect of group was not significant (F_(1,43)_ = 2.06, p = 0.159), but the interactions between hemisphere and group (F_(1,43)_ = 5.47, p = 0.024), and between hemisphere and tract (F_(2,42)_ = 7.06, p = 0.002) were significant. However, the 3-way interaction between group, tract and hemisphere did not reach significance (F_(2,42)_ = 3.01, p = 0.060). The significant effects were driven by patients having lower FA in bilateral AF/SLF (t_(43)_ = 2.24, p = 0.015; t_(43)_ = 2.20, p = 0.017, for right and left AF/SLF, respectively), with no group difference in the other tracts (p > 0.05 for all). Lastly, patients’ fornix had significantly lower FA compared with the control group (t_(43)_ = 1.87, p = 0.034). All effects remained similar when controlling for age.

### 3.3 Structure-function relationships

Within the patient group, better memory scores were associated with bilaterally larger volumes of the hippocampus (Left: Pearson’s r = 0.56, p = 0.004; Right: r = 0.46, p = 0.018), thalamus (Left: r = 0.51, p = 0.009; Right: r = 0.45, p = 0.020), caudate nucleus (Left: r = 0.44, p = 0.024; Right: r = 0.45, p = 0.021) and putamen (Left: r = 0.39 p = 0.037; Right: r = 0.45, p = 0.021), as well as whole brain white matter volume (r = 0.38, p = 0.043). See Figure 4.

**Figure 4.**
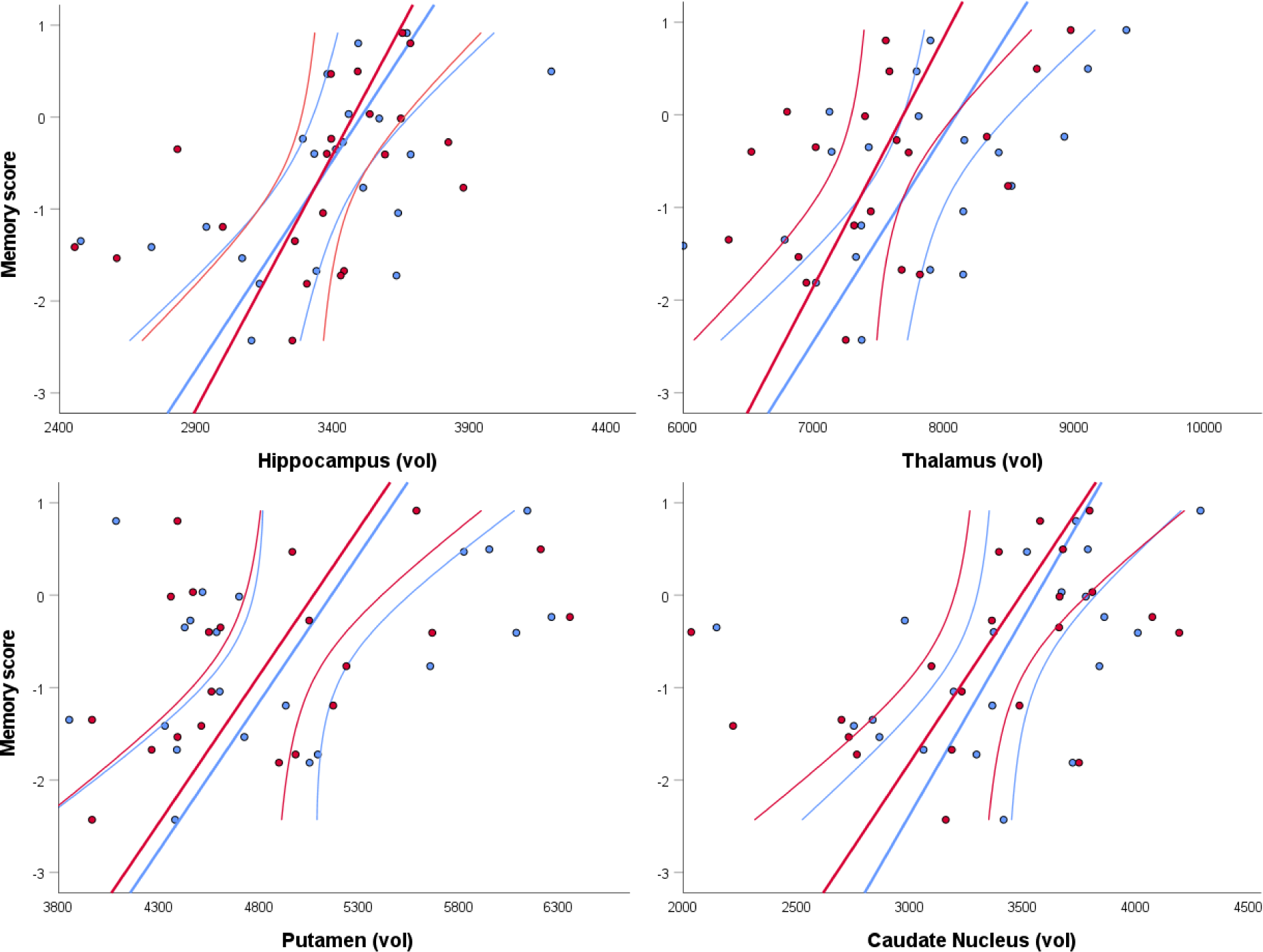
Correlations between memory scores and volumes of subcortical structures. Y axis represents memory scores derived from the PCA; x axis represents volumes of right hemispheric (red) and left hemispheric (blue) subcortical structures (mm^3^); straight lines represent linear correlations; curved lines represent 95% confidence interval of the mean.

Higher attainment scores were significantly associated with higher bilateral FA in the SCP (Left: r = 0.53, p=0.008; Right: r = 0.59, p=0.003).

Language scores were not significantly correlated with any white or grey matter parameters, (p > 0.05 for all non-significant bivariate correlations). We also note that no significant structure-function relationships were found in the group of control participants.

## 4 Discussion

We studied a rare cohort of 28 children who suffered a cardiorespiratory arrest early in life, with the aim of establishing the behavioural and neuroanatomical sequalae of this HI event. Four main findings in the patient group define the functional and anatomical outcome following paediatric CA: (i) The patient group had scores in the impaired range on memory, language and academic attainment; (ii) This was accompanied by lower values in measurements of global brain tissue volume and microstructure; (iii) Memory impairments were associated with bilaterally reduced volumes of subcortical grey matter structures, including the hippocampi, thalami, and striatum; and, (iv) Lower attainment scores were associated with reduced white matter FA in the superior cerebellar peduncle (SCP). We discuss each of these findings below.

### 4.1 Cognitive impairments and brain abnormalities

Previous studies of developmental outcome following HI have documented a range of cognitive deficits, including in the domains of memory (Cooper et al., 2015; Gadian et al., 2000; Muñoz-López et al., 2017), language (Hövels-Gürich et al., 2002; Martinez et al., 2014; Robertson and Finer, 1985), and attainment (Hövels-Gürich et al., 2002; Marlow et al., 2005), though not necessarily simultaneously. Here we provide evidence that HI of cardiorespiratory origin can lead to impairments in all three domains. This finding is in line with a study of adult CA, which suggested that memory problems rarely appear in isolation, as even when studying only those patients who were referred to a memory clinic, mild to severe motor problems, and executive function deficits were found in all 10 patients who had memory impairments (Lim et al., 2004). We note that while as a group, the patients had a deficit in all three behavioural components, inter-subject variability was large, and not all patients were impaired on all tasks (see Supplementary Table 4). In addition, variability between subtests was noted to be large as well, ranging from only one patient having impaired score on the MLU measurement, to third of patients showing impaired performance on the numerical operations subtest. Looking at the component scores, the language domain showed the lowest level of impaired performance, as only 36% of patients had a score within the impaired range (defined here as a score which is more than 1.5 standard deviations below the control group mean), followed by attainment, with 50% of patients having scores in the impaired range, and as hypothesised, the largest proportion of impaired performance was seen in the memory domain (79%).

The patient group studied here also showed global brain abnormalities, which were driven by lower whole brain white and grey matter volumes (but not CSF), and lower whole brain FA (but not MD or RD). Whole brain white matter abnormalities have been documented previously in paediatric populations with history of HI. For example, lower white matter volumes were found in neonates treated for acute respiratory failure (Cooper et al., 2015), and children born prematurely (Northam et al., 2011; Soria-Pastor et al., 2008). Widespread white matter abnormalities, quantified using diffusivity parameters (Mullen et al., 2011; Thompson et al., 2014a), and VBM (Soria-Pastor et al., 2008) have been documented in children born prematurely.

Looking at specific grey and white matter structures, we found that in our patient group, structures which are known to be vulnerable to HI in various paediatric populations, were affected. This included reduced volumes of the hippocampus (Cooper et al., 2015; Isaacs et al., 2003; Muñoz-López et al., 2017; Singh et al., 2019), thalamus (Dzieciol et al., 2017; Gadian et al., 2000; Singh et al., 2019), and caudate nucleus (Guderian et al., 2015; Singh et al., 2019), as well as lower FA values in the bilateral AF/SLF (Mullen et al., 2011), and the fornix (Singh et al., 2019). Both mouse (Stone et al., 2008) and rat models (Delcour et al., 2012) of neonatal HI showed damage to the hippocampus and fornix, with the mouse model also providing evidence that the hippocampal damage precedes the degeneration of the fornix (Stone et al., 2008). As such, our findings are in line with these previous studies, confirming that the pattern of brain damage seen following CA is similar to that seen following other HI aetiologies.

### 4.2 Memory impairment and its neural correlates

It is well known that hypoxic or ischaemic damage can cause memory impairments in both the developing (Cooper et al., 2015; Gadian et al., 1999) and adult brain (Alexander, Lafleche, Schnyer, Lim, & Verfaellie, 2011; Bunch et al., 2004; Cronberg, Lilja, Rundgren, Friberg, & Widner, 2009; Grubb, Ocarroll, Cobbe, Sirel, & Fox, 1996; Mateen et al., 2011; Moulaert, Verbunt, van Heugten, & Wade, 2009; O’Reilly, Grubb, & O’Carroll, 2003; Sauve, 1995; Sunnerhagen, Johansson, Herlitz, & Grimby, 1996; Tiainen et al., 2007), and that the hippocampus, thalamus and dorsal striatum are especially vulnerable to HI (Fatemi et al., 2009).

Early models distinguished a ‘*memory system*’ from a ‘*habit formation system*’. The memory system was thought to be dependent on the cortical-limbic-thalamic circuit with the hippocampus as an essential hub, while the habit formation system was associated with the cortico-striatal circuit (Mishkin et al., 1984). Below, we discuss each of these circuits and its potential relation to mnemonic processes, starting with the hippocampus-based circuit. The involvement of the hippocampus in cognitive memory is now widely accepted, and studies from our group confirmed the relationship between early hypoxia-induced hippocampal damage and selective memory disturbance (Cooper et al., 2015; Mishkin et al., 1998; Muñoz-López et al., 2017; Vargha-Khadem et al., 1997). Similarly, a previous study from our group provided evidence that patients with developmental amnesia following various HI events early in life, have smaller than normal mid-anterior and posterior thalamus, but only the volume of the former correlated with memory function (Dzieciol et al., 2017). Based on studies of patients with thalamic damage who suffer memory impairments, and studies of animal models, it has been suggested that in adults, the anterior thalamic nuclei are a key node for integration of information (Hwang et al., 2017; Nelson, 2021; Sweeney-Reed et al., 2021), a process which supports episodic memory formation (Aggleton, 2014). The mnemonic function of the anterior thalamic nuclei is therefore related to their structural connections to the hippocampal formation and the mammillary bodies (Grodd et al., 2020; reviewed in Sweeney-Reed et al., 2021), but also to anatomical connections beyond the hippocampal circuity (Aggleton, 2014; Nelson, 2021).

There is also evidence that the striatum and its circuity are involved in various forms of learning and habit formation (Haber, 2016; Mishkin et al., 1984). During procedural learning (also described as ‘implicit’ or ‘reflexive’) the dorsal striatum (the posterior putamen and body and tail of caudate nucleus) support the process of making a learnt response automatic (Ashby et al., 1998; Ashby and Ennis, 2006; Cohen and Frank, 2009; Ell et al., 2011; Waldschmidt and Ashby, 2011). During rule-based learning the dorsal striatum holds relevant, and filters out irrelevant information in the initial learning stages (Ashby and Ennis, 2006; Ell et al., 2006; Kovacs et al., 2021). In parallel, the hippocampus keeps track of accepted and rejected rules (Kovacs et al., 2021).

Further discussion of the role of the different dorsal striatal structures in learning is beyond the scope of the current study. However, we note that (i) it is widely accepted that the dorsal striatum is involved in various types of learning, beyond mere motor learning (Haber, 2016; Janacsek et al., 2022), supporting the findings form the current study, but (ii) only few studies have focused on the interplay between the hippocampal-based circuit and the striatal-based circuit(s) during learning processes, suggesting that, in adults, the hippocampus and caudate nucleus work together to consolidate learning [temporal (van de Ven et al., 2020) or spatial (Woolley et al., 2015)], therefore supporting memory formation.

In our study, lower integrity of both the striatum and the hippocampus was associated with compromised performance on complex memory tasks. Such performance can reflect difficulty in different stages and types of learning. Further disentanglement of the role of the caudate nucleus (head, body or tail), putamen (posterior or anterior) and hippocampus (with its multiple divisions) in learning and memory formation, will require future studies to employ (i) fine-grained behavioural experimental paradigms, and; (ii) high resolution neuroimaging, in similar developmental populations to the one studied here.

### 4.3 Impairment in attainment and its neural correlates

As the WIAT is a widely used standardised test, suitable for identifying children’s strengths and weaknesses across a range of academic areas (Smith, 2001), it is an important tool in a cohort study like ours. However, performance on each of the subtests in this battery is dependent on multiple cognitive processes, and as a result, it is difficult to determine which cognitive process primarily contributed to the documented reduced performance and to the statistical association between task performance and the integrity of the cerebellar white matter tracts. Moreover, here we calculated an attainment factor, which included a few WIAT subtests. Looking at individual subtests, word reading and spelling loaded high on this factor (>0.90) and numerical operations had slightly lower, though still substantial, factor loading (0.85). At the same time, patients were most impaired on the numerical operations subtest, with the group average significantly below the standard mean of 100, producing a large effect size. This parallels the findings of a recent study, showing that children who underwent heart transplant were impaired on the three WIAT subtests used here, with numerical operations producing the lowest group mean (Gold et al., 2020). While numerical operations are not often studied in developmental populations with history of HI, our study still replicates findings from some studies where difficulties with numerical operations were found (Isaacs et al., 2001; Taylor et al., 2011).

In addition, we found that attainment scores were associated with the integrity of the SCP bilaterally, which is the main efferent white matter bundle of the cerebellum. Previous studies of children with cerebellar tumour resections have associated low SCP integrity [measured either using lesion symptom mapping (Albazron et al., 2019; Grosse et al., 2021); or FA (Law et al., 2017, 2011; Rueckriegel et al., 2015)] with general cognitive impairment (Albazron et al., 2019; Grosse et al., 2021; Rueckriegel et al., 2015), or more specifically, with impairments of working memory (Law et al., 2017, 2011). Together, these studies provide growing evidence for the relation between damage to the cerebellar efferent pathways and cognitive function in paediatric populations. Our study is the first to demonstrate this relation in a population with history of HI.

### 4.4 Impairments in language processing

The patient group in our study had impaired scores on the language component. At the same time, we also note that the group impairment was subtle, with most patients performing within the normal range (see Figure 1 and Supplementary Table 4). In addition, we did not find an association between any of the grey or white matter structures studied here, and the language scores. This might be a consequence of the aggregated nature of the score, i.e. it combines measurements of language comprehension and production, which are likely to depend on partially different neural network (Giglio et al., 2022). In contrast to our study results, previous reports of developmental outcome following neonatal HI have successfully associated the occurrence and severity of speech and language disorders with the level of damage to the basal ganglia (Martinez-Biarge et al., 2012b, 2012a, 2010). In another study, verbal IQ was associated with watershed white matter damage (Steinman et al., 2009). In these previous studies patients had severe deficits and substantial damage which could be seen on clinical MRI images. In our study, severe visible damage was documented in only a handful of patients (see Supplementary Table 3) and cognitive impairments were much more subtle, which together might explain the difficulty in associating neural mechanisms and behavioural outcome. Future studies are needed to measure more fine-grained cognitive mechanisms involved in language processing, which in turn might be more easily related to underlying neural mechanisms.

### 4.5 Study caveats and future work

Studying such a unique cohort of patients poses many challenges. First, we did not identify a single medical variable, or combination of medical variables, that could explain patients’ cognitive outcome. One possible explanation is that the variables recorded in this study were not sensitive enough to capture the severity of the HI event. For example, downtime (total time of cardiorespiratory arrest) is usually documented by the clinical staff present during the event, after resuscitation is achieved and the patient is stabilised. When the cardiac arrest occurs outside hospital, downtime measurement will be based on evidence given by the person present during the event. In both cases, the documented downtime is an approximation, and as such might not be accurate enough to statistically explain long-term cognitive outcome. In line with our study, previous investigations of adult populations have also failed to identify medical variables that predict functional outcome (Mateen et al., 2011; Moulaert et al., 2009; O’Reilly et al., 2003; Sunnerhagen et al., 1996; Tiainen et al., 2007). Similarly, a recent systematic review (Moulaert et al., 2009) concluded that resuscitation variables and co-morbidities are not conclusively related to cognitive outcome. However, they have suggested that being awake at admission and a short duration of coma are related to better long-term cognitive functioning. Likewise, Grubb et al., (1996) found that, among adults, duration of cardiac arrest was a reliable predictor of memory impairment. Similarly, a study of neurodevelopmental status following neonatal TGA operation found that severe preoperative acidosis and hypoxia were associated with poor cognitive outcome (Hövels-Gürich et al., 2002). Notably, this was defined based on more subtle measurements compared to those obtained in our study (e.g., pH value <7.2 in umbilical venous blood), which in turn might contribute to better outcome prediction.

Second, individuals in our patient cohort suffered their CA as a result of varied aetiologies, comorbidities (whether direct outcome from the main diagnosis or not), and the presence, absence, and type of ongoing medical problems. These factors were not addressed in our statistical analyses, as the variability made it impossible to code them in a way that would allow for quantitative analysis. We suggest that future studies could look at the overall severity of the medical condition, as defined by the combination of a few theoretically relevant medical variables. Such aggregated score might be informative when it comes to predicting patients’ outcome.

Third, we did not measure non-medical variables which might influence the child’s performance on cognitive and achievement tests, such as school absence, parental support, or help given outside the school environment. Some of these variables can be estimated using other available data. For example, school absence estimation can be based on out-patient appointments in our hospital, however, for many children this will not reflect actual absence, either because their main care service is local (i.e., not in our hospital), or because they might miss out on school days due to different, though related reasons. The influence of such parameters on the child’s cognitive outcome should be evaluated separately and systematically.

Lastly, future studies should evaluate the implications of the behavioural deficits documented here on the quality of life of the children and/or their carers. This is especially important in light of previous findings in adults, showing that memory and other cognitive impairments resulting from hypoxic brain damage affect independence of patients, and the quality of life of both patient and their carers (Pusswald, Fertl, Faltl, & Auff, 2000).

## 5 Conclusions

Children with a history of early cardiorespiratory arrest are an under-studied group of patients. In this cohort study we have found that later in life, those children show substantial deficits in the areas of memory, language and academic attainment. These domains require further investigation and should be addressed clinically in children with a similar medical history as the ones investigated here. Impairments in memory were highlighted in the past extensively, and currently it is well established that hypoxia-ischaemia can lead to such outcome, in both adults and children. However, the deficit in language and attainment require further attention. As such, our findings of cognitive impairments across various domains, together with global brain abnormalities which are driven by damage to specific grey and white matter structures, add to the current understanding of long-term outcome following paediatric HI.

## 6 Acknowledgements

We thank Eva Zita Patai, Georgia Pitts, Steve Russon, and the team at Great Ormond Street Hospital Cardiac Intensive Care Unit (CICU), for helping with recruitment and assessment of participants. This research was supported by the NIHR Great Ormond Street Hospital Biomedical Research Centre. We also thank the participants and their families for taking part in this study.

## 8 References

Aggleton, J.P., 2014. Looking beyond the hippocampus: old and new neurological targets for understanding memory disorders. Proc. R. Soc. B Biol. Sci. 281. https://doi.org/10.1098/RSPB.2014.0565

Albazron, F.M., Bruss, J., Jones, R.M., Yock, T.I., Pulsifer, M.B., Cohen, A.L., Nopoulos, P.C., Abrams, A.N., Sato, M., Boes, A.D., 2019. Pediatric postoperative cerebellar cognitive affective syndrome follows outflow pathway lesions. Neurology 93, E1561–E1571. https://doi.org/10.1212/WNL.0000000000008326

Alexander, M.P., Lafleche, G., Schnyer, D., Lim, C., Verfaellie, M., 2011. Cognitive and Functional Outcome After Out of Hospital Cardiac Arrest. J. Int. Neuropsychol. Soc. 17, 364–368. https://doi.org/10.1017/s1355617710001633

Ashburner, J., 2007. A fast diffeomorphic image registration algorithm. Neuroimage 38, 95–113. https://doi.org/10.1016/j.neuroimage.2007.07.007

Ashburner, J., Friston, K.J., 2005. Unified segmentation. Neuroimage 26, 839–851. https://doi.org/10.1016/j.neuroimage.2005.02.018

Ashby, F.G., Alfonso-Reese, L.A., Turken, A.U., Waldron, E.M., 1998. A Neuropsychological Theory of Multiple Systems in Category Learning. Psychol. Rev. 105, 442–481. https://doi.org/10.1037/0033-295X.105.3.442

Ashby, F.G., Ennis, J.M., 2006. The Role of the Basal Ganglia in Category Learning. Psychol. Learn. Motiv. - Adv. Res. Theory. https://doi.org/10.1016/S0079-7421(06)46001-1

Benjamini, Y., Hochberg, Y., 1995. Controlling the false discovery rate: a practical and powerful approach to multiple testing. J. R. Stat. Soc. 57, 289–300. https://doi.org/10.2307/2346101

Bertini, G., Giglioli, C., Giovannini, F., Bartoletti, A., Cricelli, F., Margheri, M., Russo, L., Taddei, T., Taiti, A., 1990. Neuropsychological outcome of survivors of out-of-hospital cardiac arrest. J. Emerg. Med. 8, 407–412. https://doi.org/10.1016/0736-4679(90)90166-S

Bishop, D.V.M., 2004. Expression, Reception and Recall of Narrative Instrument.

Buck, S., Bastos, F., Baldeweg, T., Vargha-Khadem, F., 2021. The Pair Test: A computerised measure of learning and memory. Behav. Res. Methods 53, 928–942. https://doi.org/10.3758/S13428-020-01470-9/TABLES/9

Bunch, T.J., White, R.D., Smith, G.E., Hodge, D.O., Gersh, B.J., Hammill, S.C., Shen, W.K., Packer, D.L., 2004. Long-term subjective memory function in ventricular fibrillation out-of-hospital cardiac arrest survivors resuscitated by early defibrillation. Resuscitation 60, 189–195. https://doi.org/10.1016/j.resuscitation.2003.09.010

Cohen, M., 1997. Children’s Memory Scale.

Cohen, M.X., Frank, M.J., 2009. Neurocomputational models of basal ganglia function in learning, memory and choice. Behav. Brain Res. 199, 141–156. https://doi.org/10.1016/J.BBR.2008.09.029

Cooper, J.M., Gadian, D.G., Jentschke, S., Goldman, A., Munoz, M., Pitts, G., Banks, T., Chong, W.K., Hoskote, A., Deanfield, J., Baldeweg, T., de Haan, M., Mishkin, M., Vargha-Khadem, F., 2015. Neonatal Hypoxia, Hippocampal Atrophy, and Memory Impairment: Evidence of a Causal Sequence. Cereb. Cortex 25, 1469–76. https://doi.org/10.1093/cercor/bht332

Corbetta, M., Ramsey, L., Callejas, A., Baldassarre, A., Hacker, C.D., Siegel, J.S., Astafiev, S. V., Rengachary, J., Zinn, K., Lang, C.E., Connor, L.T., Fucetola, R., Strube, M., Carter, A.R., Shulman, G.L., 2015. Common behavioral clusters and subcortical anatomy in stroke. Neuron 85, 927. https://doi.org/10.1016/J.NEURON.2015.02.027

Cowan, F., Rutherford, M., Groenendaal, F., Eken, P., Mercuri, E., Bydder, G.M., Meiners, L.C., Dubowitz, L.M.S., de Vries, L.S., 2003. Origin and timing of brain lesions in term infants with neonatal encephalopathy. Lancet (London, England) 361, 736–42. https://doi.org/10.1016/S0140-6736(03)12658-X

Cronberg, T., Lilja, G., Rundgren, M., Friberg, H., Widner, H., 2009. Long-term neurological outcome after cardiac arrest and therapeutic hypothermia. Resuscitation 80, 1119– 1123. https://doi.org/10.1016/j.resuscitation.2009.06.021

Delcour, M., Russier, M., Amin, M., Baud, O., Paban, V., Barbe, M.F., Coq, J.O., 2012. Impact of prenatal ischemia on behavior, cognitive abilities and neuroanatomy in adult rats with white matter damage. Behav. Brain Res. 232, 233–244. https://doi.org/10.1016/J.BBR.2012.03.029

Dick, A.S., Bernal, B., Tremblay, P., 2014. The language connectome: new pathways, new concepts. Neuroscientist 20, 453–467. https://doi.org/10.1177/1073858413513502

Dogan, E., Gungor, A., Dogulu, F., Türe, U., 2022. The historical evolution of the fornix and its terminology: a review. Neurosurg. Rev. 45, 979–988. https://doi.org/10.1007/S10143-021-01635-W

Drysdale, E.E., Grubb, N.R., Fox, K.A.A., O’Carroll, R.E., 2000. Chronicity of memory impairment in long-term out-of-hospital cardiac arrest survivors. Resuscitation 47, 27–32. https://doi.org/10.1016/S0300-9572(00)00194-5

Dzieciol, A.M., Bachevalier, J., Saleem, K.S., Gadian, D.G., Saunders, R., Chong, W.K.K.K., Banks, T., Mishkin, M., Vargha-Khadem, F., 2017. Hippocampal and diencephalic pathology in developmental amnesia. Cortex 86, 33–44. https://doi.org/10.1016/j.cortex.2016.09.016

Ell, S., Helie, S., Hutchinson, S., 2011. Contributions of the putamen to cognitive function, in: Costa, A., Villalba, E. (Eds.), Horizon in Neuroscience. Nova Science Publishers, Hauppauge, NY, pp. 29–52.

Ell, S.W., Marchant, N.L., Ivry, R.B., 2006. Focal putamen lesions impair learning in rule-based, but not information-integration categorization tasks. Neuropsychologia 44, 1737–1751. https://doi.org/10.1016/j.neuropsychologia.2006.03.018

Engdahl, J., Axelsson, Å., Bång, A., Karlson, B.W., Herlitz, J., 2003. The epidemiology of cardiac arrest in children and young adults. Resuscitation 58, 131–138. https://doi.org/10.1016/S0300-9572(03)00108-4

Fatemi, A., Wilson, M.A., Johnston, M. V., 2009. Hypoxic-Ischemic Encephalopathy in the Term Infant. Clin. Perinatol. 36, 835–58, vii. https://doi.org/10.1016/j.clp.2009.07.011

Friederici, A.D., 2015. White-matter pathways for speech and language processing. Handb. Clin. Neurol. 129, 177–186. https://doi.org/10.1016/B978-0-444-62630-1.00010-X

Gadian, D.G., Aicardi, J., Watkins, K.E., Porter, D.A., Misbkin, M., Vargha-Khadem, F., 1999. Developmental amnesia associated with early hypoxic-ischemic injury. Soc. Neurosci. Abstr. 25, 894.

Gadian, D.G., Aicardi, J., Watkins, K.E., Porter, D.A., Mishkin, M., Vargha-Khadem, F., 2000. Developmental amnesia associated with early hypoxic-ischaemic injury. Brain 123, 499–507. https://doi.org/10.1093/brain/123.3.499

Giglio, L., Ostarek, M., Weber, K., Hagoort, P., 2022. Commonalities and Asymmetries in the Neurobiological Infrastructure for Language Production and Comprehension. Cereb. Cortex 32, 1405–1418. https://doi.org/10.1093/CERCOR/BHAB287

Gold, A., Bondi, B.C., Ashkanase, J., Dipchand, A.I., 2020. Early school-age cognitive performance post–pediatric heart transplantation. Pediatr. Transplant. 24, e13832. https://doi.org/10.1111/PETR.13832

Grodd, W., Kumar, V.J., Schüz, A., Lindig, T., Scheffler, K., 2020. The anterior and medial thalamic nuclei and the human limbic system: tracing the structural connectivity using diffusion-weighted imaging. Sci. Rep. 10, 1–25. https://doi.org/10.1038/s41598-020-67770-4

Grosse, F., Rueckriegel, S.M., Thomale, U.W., Hernáiz Driever, P., 2021. Mapping of long-term cognitive and motor deficits in pediatric cerebellar brain tumor survivors into a cerebellar white matter atlas. Child’s Nerv. Syst. 37, 2787–2797. https://doi.org/10.1007/S00381-021-05244-2/FIGURES/3

Grubb, N.R., Fox, K.A.A., Smith, K., Best, J., Blane, A., Ebmeier, K.P., Glabus, M.F., O’Carroll, R.E., 2000. Memory impairment in out-of-hospital cardiac arrest survivors is associated with global reduction in brain volume, not focal hippocampal injury. Stroke 31, 1509– 1514.

Grubb, N.R., Ocarroll, R., Cobbe, S.M., Sirel, J., Fox, K.A.A., 1996. Chronic memory impairment after cardiac arrest outside hospital. Br. Med. J. 313, 143–146.

Guderian, S., Dzieciol, A.M., Gadian, D.G., Jentschke, S., Doeller, C.F., Burgess, N., Mishkin, M., Vargha-Khadem, F., 2015. Hippocampal Volume Reduction in Humans Predicts Impaired Allocentric Spatial Memory in Virtual-Reality Navigation. J. Neurosci. 35, 14123–14131. https://doi.org/10.1523/JNEUROSCI.0801-15.2015

Haber, S.N., 2016. Corticostriatal circuitry. Dialogues Clin. Neurosci. 18, 7–21. https://doi.org/10.1007/978-1-4614-6434-1_135-1

Holmberg, M.J., Ross, C.E., Fitzmaurice, G.M., Chan, P.S., Duval-Arnould, J., Grossestreuer, A. V., Yankama, T., Donnino, M.W., Andersen, L.W., 2019. Annual Incidence of Adult and Pediatric In-Hospital Cardiac Arrest in the United States. Circ. Cardiovasc. Qual. Outcomes 12. https://doi.org/10.1161/CIRCOUTCOMES.119.005580

Hövels-Gürich, H.H., Seghaye, M.-C.C., Schnitker, R., Wiesner, M., Huber, W., Minkenberg, R., Kotlarek, F., Messmer, B.J., Von Bernuth, G., 2002. Long-term neurodevelopmental outcomes in school-aged children after neonatal arterial switch operation. J. Thorac. Cardiovasc. Surg. 124, 448–458. https://doi.org/10.1067/mtc.2002.122307

Hwang, K., Bertolero, M.A., Liu, W.B., D’Esposito, M., 2017. The human thalamus is an integrative hub for functional brain networks. J. Neurosci. 37, 5594–5607. https://doi.org/10.1523/JNEUROSCI.0067-17.2017

Ichord, R., Silverstein, F.S., Slomine, B.S., Telford, R., Christensen, J., Holubkov, R., Dean, J.M., Moler, F.W., THAPCA Trial Group, 2018. Neurologic outcomes in pediatric cardiac arrest survivors enrolled in the THAPCA trials. Neurology 91, e123–e131. https://doi.org/10.1212/WNL.0000000000005773

Isaacs, E.B., Edmonds, C.J., Lucas, A., Gadian, D.G., 2001. Calculation difficulties in children of very low birthweight: a neural correlate. Brain 124, 1701–7. https://doi.org/10.1093/brain/124.9.1701

Isaacs, E.B., Vargha-Khadem, F., Watkins, K.E., Lucas, A., Mishkin, M., Gadian, D.G., 2003. Developmental amnesia and its relationship to degree of hippocampal atrophy. Proc. Natl. Acad. Sci. U. S. A. 100, 13060–13063. https://doi.org/10.1073/pnas.1233825100

Janacsek, K., Evans, T.M., Kiss, M., Shah, L., Blumenfeld, H., Ullman, M.T., 2022. Subcortical Cognition: The Fruit Below the Rind. Annu. Rev. Neurosci. 45, 361–386. https://doi.org/10.1146/ANNUREV-NEURO-110920-013544

Kovacs, P., Hélie, S., Tran, A.N., Ashby, F.G., 2021. A neurocomputational theory of how rule-guided behaviors become automatic. Psychol. Rev. 128, 488–508. https://doi.org/10.1037/rev0000271

Law, N., Bouffet, E., Laughlin, S., Laperriere, N., Brière, M.E., Strother, D., McConnell, D., Hukin, J., Fryer, C., Rockel, C., Dickson, J., Mabbott, D., 2011. Cerebello-thalamo-cerebral connections in pediatric brain tumor patients: impact on working memory. Neuroimage 56, 2238–2248. https://doi.org/10.1016/J.NEUROIMAGE.2011.03.065

Law, N., Smith, M. Lou, Greenberg, M., Bouffet, E., Taylor, M.D., Laughlin, S., Malkin, D., Liu, F., Moxon-Emre, I., Scantlebury, N., Mabbott, D., 2017. Executive function in paediatric medulloblastoma: The role of cerebrocerebellar connections 11, 174–200. https://doi.org/10.1111/JNP.12082

Lim, C., Alexander, M.P., LaFleche, G., Schnyer, D.M., Verfaellie, M., 2004. The neurological and cognitive sequelae of cardiac arrest. Neurology 63, 1774–1778.

Manole, M.D., Kochanek, P.M., Fink, E.L., Clark, R.S., 2009. Postcardiac arrest syndrome: Focus on the brain. Curr. Opin. Pediatr. https://doi.org/10.1097/MOP.0b013e328331e873

Marlow, N., Rose, A.S., Rands, C.E., Draper, E.S., 2005. Neuropsychological and educational problems at school age associated with neonatal encephalopathy. Arch. Dis. Child. Fetal Neonatal Ed. 90, F380–F387. https://doi.org/10.1136/adc.2004.067520

Martinez-Biarge, M., Bregant, T., Wusthoff, C.J., Chew, A.T.M., Diez-Sebastian, J., Rutherford, M.A., Cowan, F.M., 2012a. White Matter and Cortical Injury in Hypoxic-Ischemic Encephalopathy: Antecedent Factors and 2-Year Outcome. J. Pediatr. 161, 799–807. https://doi.org/10.1016/j.jpeds.2012.04.054

Martinez-Biarge, M., Diez-Sebastian, J., Rutherford, M.A., Cowan, F.M., 2010. Outcomes after central grey matter injury in term perinatal hypoxic-ischaemic encephalopathy. Early Hum. Dev. 86, 675–682. https://doi.org/10.1016/j.earlhumdev.2010.08.013

Martinez-Biarge, M., Diez-Sebastian, J., Wusthoff, C.J., Lawrence, S., Aloysius, A., Rutherford, M.A., Cowan, F.M., 2012b. Feeding and communication impairments in infants with central grey matter lesions following perinatal hypoxic-ischaemic injury. Eur. J. Paediatr. Neurol. 16, 688–96. https://doi.org/10.1016/j.ejpn.2012.05.001

Martinez, C., Carneiro, L., Vernier, L., Cesa, C., Guardiola, A., Vidor, D., 2014. Language in children with neonatal hypoxic-ischemic encephalopathy. Int. Arch. Otorhinolaryngol. 18, 255–9. https://doi.org/10.1055/s-0034-1366976

Maryniak, A., Bielawska, A., Walczak, F., Szumowski, Ł., Bieganowska, K., Rękawek, J., Paszke, M., Szymaniak, E., Knecht, M., 2008. Long-term cognitive outcome in teenage survivors of arrhythmic cardiac arrest. Resuscitation 77, 46–50. https://doi.org/10.1016/j.resuscitation.2007.10.024

Mateen, F.J., Josephs, K.A., Trenerry, M.R., Felmlee-Devine, M.D., Weaver, A.L., Carone, M., White, R.D., 2011. Long-term cognitive outcomes following out-of-hospital cardiac arrest A population-based study. Neurology 77, 1438–1445. https://doi.org/10.1212/WNL.0b013e318232ab33

Matejko, A.A., Ansari, D., 2015. Drawing connections between white matter and numerical and mathematical cognition: A literature review. Neurosci. Biobehav. Rev. 48, 35–52. https://doi.org/10.1016/j.neubiorev.2014.11.006

Meert, K., Telford, R., Holubkov, R., Slomine, B.S., Christensen, J.R., Berger, J., Ofori-Amanfo, G., Newth, C.J.L., Dean, J.M., Moler, F.W., 2018. Paediatric in-hospital cardiac arrest: Factors associated with survival and neurobehavioural outcome one year later. Resuscitation 124, 96–105. https://doi.org/10.1016/J.RESUSCITATION.2018.01.013

Meert, K.L., Telford, R., Holubkov, R., Slomine, B.S., Christensen, J.R., Dean, J.M., Moler, F.W., 2016. Pediatric Out-of-Hospital Cardiac Arrest Characteristics and Their Association With Survival and Neurobehavioral Outcome. Pediatr. Crit. Care Med. 17, e543–e550. https://doi.org/10.1097/PCC.0000000000000969

Mishkin, M., Malamut, B., Bachevalier, J., 1984. Memories and habits: Two neural systems. Guilford, New York.

Mishkin, M., Vargha-Khadem, F., Gadian, D.G., 1998. Amnesia and the organization of the hippocampal system. Hippocampus 8, 212–216. https://doi.org/10.1002/(sici)1098-1063(1998)8:3<212::aid-hipo4>3.0.co;2-l

Moeller, K., Willmes, K., Klein, E., 2015. A review on functional and structural brain connectivity in numerical cognition. Front. Hum. Neurosci. 9, 227. https://doi.org/10.3389/fnhum.2015.00227

Moler, F.W., Silverstein, F.S., Holubkov, R., Slomine, B.S., Christensen, J.R., Nadkarni, V.M., Meert, K.L., Browning, B., Pemberton, V.L., Page, K., Gildea, M.R., Scholefield, B.R., Shankaran, S., Hutchison, J.S., Berger, J.T., Ofori-Amanfo, G., Newth, C.J.L., Topjian, A., Bennett, K.S., Koch, J.D., Pham, N., Chanani, N.K., Pineda, J.A., Harrison, R., Dalton, H.J., Alten, J., Schleien, C.L., Goodman, D.M., Zimmerman, J.J., Bhalala, U.S., Schwarz, A.J., Porter, M.B., Shah, S., Fink, E.L., McQuillen, P., Wu, T., Skellett, S., Thomas, N.J., Nowak, J.E., Baines, P.B., Pappachan, J., Mathur, M., Lloyd, E., van der Jagt, E.W., Dobyns, E.L., Meyer, M.T., Sanders, R.C., Clark, A.E., Dean, J.M., 2017. Therapeutic Hypothermia after In-Hospital Cardiac Arrest in Children. N. Engl. J. Med. 376, 318–329. https://doi.org/10.1056/NEJMOA1610493

Moler, F.W., Silverstein, F.S., Holubkov, R., Slomine, B.S., Christensen, J.R., Nadkarni, V.M., Meert, K.L., Clark, A.E., Browning, B., Pemberton, V.L., Page, K., Shankaran, S., Hutchison, J.S., Newth, C.J.L., Bennett, K.S., Berger, J.T., Topjian, A., Pineda, J.A., Koch, J.D., Schleien, C.L., Dalton, H.J., Ofori-Amanfo, G., Goodman, D.M., Fink, E.L., McQuillen, P., Zimmerman, J.J., Thomas, N.J., van der Jagt, E.W., Porter, M.B., Meyer, M.T., Harrison, R., Pham, N., Schwarz, A.J., Nowak, J.E., Alten, J., Wheeler, D.S., Bhalala, U.S., Lidsky, K., Lloyd, E., Mathur, M., Shah, S., Wu, T., Theodorou, A.A., Sanders, R.C., Dean, J.M., 2015. Therapeutic Hypothermia after Out-of-Hospital Cardiac Arrest in Children. N. Engl. J. Med. 372, 1898–1908. https://doi.org/10.1056/nejmoa1411480

Mori, S., Wakana, S., van Zijl, P.C.M., Nagae-Poetscher, L.M., 2005. MRI atlas of human white matter, 1st ed. Elsevier.

Moulaert, V.R.M.P., Verbunt, J.A., van Heugten, C.M., Wade, D.T., 2009. Cognitive impairments in survivors of out-of-hospital cardiac arrest: A systematic review. Resuscitation 80, 297–305. https://doi.org/10.1016/j.resuscitation.2008.10.034

Mullen, K.M., Vohr, B.R., Katz, K.H., Schneider, K.C., Lacadie, C., Hampson, M., Makuch, R.W., Reiss, A.L., Constable, R.T., Ment, L.R., 2011. Preterm birth results in alterations in neural connectivity at age 16 years. Neuroimage 54, 2563–70. https://doi.org/10.1016/j.neuroimage.2010.11.019

Muñoz-López, M., Hoskote, A., Chadwick, M.J., Dzieciol, A.M., Gadian, D.G., Chong, K., Banks, T., de Haan, M., Baldeweg, T., Mishkin, M., Vargha-Khadem, F., 2017. Hippocampal damage and memory impairment in congenital cyanotic heart disease. Hippocampus 27, 417–424. https://doi.org/10.1002/hipo.22700

Nelson, A.J.D., 2021. The anterior thalamic nuclei and cognition: A role beyond space? Neurosci. Biobehav. Rev. https://doi.org/10.1016/j.neubiorev.2021.02.047

Northam, G.B., Liégeois, F., Chong, W.K., Wyatt, J.S., Baldeweg, T., 2011. Total brain white matter is a major determinant of IQ in adolescents born preterm. Ann. Neurol. 69, 702–11. https://doi.org/10.1002/ana.22263

Nunes, B., Pais, J., Garcia, R., Magalhaes, Z., Granja, C., Silva, M.C., 2003. Cardiac arrest: long-term cognitive and imaging analysis. Resuscitation 57, 287–297. https://doi.org/10.1016/s0300-9572(03)00033-9

O’Reilly, S.M., Grubb, N.R., O’Carroll, R.E., 2003. In-hospital cardiac arrest leads to chronic memory impairment. Resuscitation 58, 73–79. https://doi.org/10.1016/s0300-9572(03)00114-x

Oldfield, R.C., 1971. The assessment and analysis of handedness: The Edinburgh inventory. Neuropsychologia 9, 97–113. https://doi.org/10.1016/0028-3932(71)90067-4

Patenaude, B., Smith, S.M., Kennedy, D.N., Jenkinson, M., 2011. A Bayesian model of shape and appearance for subcortical brain segmentation. Neuroimage 56, 907–922. https://doi.org/10.1016/j.neuroimage.2011.02.046

Pusswald, G., Fertl, E., Faltl, M., Auff, E., 2000. Neurological rehabilitation of severely disabled cardiac arrest survivors. Part II. Life situation of patients and families after treatment. Resuscitation 47, 241–248. https://doi.org/10.1016/s0300-9572(00)00240-9

Ramsey, L.E., Siegel, J.S., Lang, C.E., Strube, M., Shulman, G.L., Corbetta, M., 2017. Behavioural clusters and predictors of performance during recovery from stroke. Nat. Hum. Behav. 1. https://doi.org/10.1038/S41562-016-0038

Robertson, C.M., Finer, N.N., 1985. Term infants with hypoxic-ischemic encephalopathy: outcome at 3.5 years. Dev. Med. Child Neurol. 27, 473–84.

Rueckriegel, S.M., Bruhn, H., Thomale, U.W., Hernáiz Driever, P., 2015. Cerebral white matter fractional anisotropy and tract volume as measured by MR imaging are associated with impaired cognitive and motor function in pediatric posterior fossa tumor survivors. Pediatr. Blood Cancer 62, 1252–1258. https://doi.org/10.1002/PBC.25485

Salvalaggio, A., de Filippo De Grazia, M., Zorzi, M., de Schotten, M.T., Corbetta, M., 2020. Post-stroke deficit prediction from lesion and indirect structural and functional disconnection. Brain 143, 2173–2188. https://doi.org/10.1093/BRAIN/AWAA156

Sauve, M.J., 1995. Long-term physical functioning and psychosocial adjustment in survivors of sudden cardiac death. Hear. Lung 24, 133–144. https://doi.org/10.1016/s0147-9563(05)80008-1

Scholefield, B.R., Silverstein, F.S., Telford, R., Holubkov, R., Slomine, B.S., Meert, K.L., Christensen, J.R., Nadkarni, V.M., Dean, J.M., Moler, F.W., 2018. Therapeutic hypothermia after paediatric cardiac arrest: Pooled randomized controlled trials. Resuscitation 133, 101–107. https://doi.org/10.1016/J.RESUSCITATION.2018.09.011

Singh, S., Roy, B., Pike, N., Daniel, E., Ehlert, L., Lewis, A.B., Halnon, N., Woo, M.A., Kumar, R., 2019. Altered brain diffusion tensor imaging indices in adolescents with the Fontan palliation. Neuroradiology 61, 811–824. https://doi.org/10.1007/s00234-019-02208-x

Slomine, B.S., Silverstein, F.S., Christensen, J.R., Holubkov, R., Page, K., Michael Dean, J., Moler, F.W., 2016. Neurobehavioral outcomes in children after out-of-hospital cardiac arrest. Pediatrics 137. https://doi.org/10.1542/peds.2015-3412

Slomine, B.S., Silverstein, F.S., Christensen, J.R., Holubkov, R., Telford, R., Dean, J.M., Moler, F.W., 2018a. Neurobehavioural outcomes in children after In-Hospital cardiac arrest. Resuscitation 124, 80–89. https://doi.org/10.1016/J.RESUSCITATION.2018.01.002

Slomine, B.S., Silverstein, F.S., Christensen, J.R., Page, K., Holubkov, R., Dean, J.M., Moler, F.W., 2018b. Neuropsychological Outcomes of Children 1 Year After Pediatric Cardiac Arrest: Secondary Analysis of 2 Randomized Clinical Trials. JAMA Neurol. 75, 1502–1510. https://doi.org/10.1001/JAMANEUROL.2018.2628

Smith, D.R., 2001. Wechsler Individual Achievement Test, in: Handbook of Psychoeducational Assessment. Elsevier Science & Technology, San Diego, pp. 169–193. https://doi.org/10.1016/B978-012058570-0/50008-2

Smith, S.M., Jenkinson, M., Woolrich, M.W., Beckmann, C.F., Behrens, T.E.J., Johansen-Berg, H., Bannister, P.R., De Luca, M., Drobnjak, I., Flitney, D.E., Niazy, R.K., Saunders, J., Vickers, J., Zhang, Y., De Stefano, N., Brady, J.M., Matthews, P.M., 2004. Advances in functional and structural MR image analysis and implementation as FSL. Neuroimage 23, S208–S219. https://doi.org/10.1016/j.neuroimage.2004.07.051

Soria-Pastor, S., Gimenez, M., Narberhaus, A., Falcon, C., Botet, F., Bargallo, N., Mercader, J.M., Junque, C., 2008. Patterns of cerebral white matter damage and cognitive impairment in adolescents born very preterm. Int. J. Dev. Neurosci. 26, 647–654. https://doi.org/10.1016/j.ijdevneu.2008.08.001

Steinman, K.J., Gorno-Tempini, M.L., Glidden, D. V., Kramer, J.H., Miller, S.P., Barkovich, A.J., Ferriero, D.M., 2009. Neonatal Watershed Brain Injury on Magnetic Resonance Imaging Correlates With Verbal IQ at 4 Years. Pediatrics 123, 1025–1030. https://doi.org/10.1542/peds.2008-1203

Stone, B.S., Zhang, J., Mack, D.W., Mori, S., Martin, L.J., Northington, F.J., 2008. Delayed neural network degeneration after neonatal hypoxia-ischemia. Ann. Neurol. 64, 535–546. https://doi.org/10.1002/ANA.21517

Sunderland, A., Harris, J.E., Baddeley, A.D., 1983. Do laboratory tests predict everyday memory? A neuropsychological study. J. Verbal Learning Verbal Behav. 22, 341–357. https://doi.org/10.1016/S0022-5371(83)90229-3

Sunnerhagen, K.S., Johansson, O., Herlitz, J., Grimby, G., 1996. Life after cardiac arrest; A retrospective study. Resuscitation 31, 135–140. https://doi.org/10.1016/0300-9572(95)00903-5

Sweeney-Reed, C.M., Buentjen, L., Voges, J., Schmitt, F.C., Zaehle, T., Kam, J.W.Y., Kaufmann, J., Heinze, H.J., Hinrichs, H., Knight, R.T., Rugg, M.D., 2021. The role of the anterior nuclei of the thalamus in human memory processing. Neurosci. Biobehav. Rev. 126, 146–158. https://doi.org/10.1016/J.NEUBIOREV.2021.02.046

Taylor, H.G., Klein, N., Anselmo, M.G., Minich, N., Espy, K.A., Hack, M., 2011. Learning Problems in Kindergarten Students With Extremely Preterm Birth. Arch. Pediatr. Adolesc. Med. 165, 819. https://doi.org/10.1001/archpediatrics.2011.137

Thompson, D.K., Lee, K.J., Egan, G.F., Warfield, S.K., Doyle, L.W., Anderson, P.J., Inder, T.E., 2014a. Regional white matter microstructure in very preterm infants: predictors and 7 year outcomes. Cortex 52, 60–74. https://doi.org/10.1016/j.cortex.2013.11.010

Thompson, D.K., Omizzolo, C., Adamson, C., Lee, K.J., Stargatt, R., Egan, G.F., Doyle, L.W., Inder, T.E., Anderson, P.J., 2014b. Longitudinal growth and morphology of the hippocampus through childhood: Impact of prematurity and implications for memory and learning. Hum. Brain Mapp. 35, 4129–39. https://doi.org/10.1002/hbm.22464

Tiainen, M., Poutiainen, E., Kovala, T., Takkunen, O., Happola, O., Roine, R.O., 2007. Cognitive and neurophysiological outcome of cardiac arrest survivors treated with therapeutic hypothermia. Stroke 38, 2303–2308. https://doi.org/10.1161/strokeaha.107.483867

Tress, E.E., Kochanek, P.M., Saladino, R.A., Manole, M.D., 2010. Cardiac arrest in children. J. Emerg. Trauma. Shock 3, 267–72. https://doi.org/10.4103/0974-2700.66528

Tsao, C.W., Aday, A.W., Almarzooq, Z.I., Alonso, A., Beaton, A.Z., Bittencourt, M.S., Boehme, A.K., Buxton, A.E., Carson, A.P., Commodore-Mensah, Y., Elkind, M.S.V., Evenson, K.R., Eze-Nliam, C., Ferguson, J.F., Generoso, G., Ho, J.E., Kalani, R., Khan, S.S., Kissela, B.M., Knutson, K.L., Levine, D.A., Lewis, T.T., Liu, J., Loop, M.S., Ma, J., Mussolino, M.E., Navaneethan, S.D., Perak, A.M., Poudel, R., Rezk-Hanna, M., Roth, G.A., Schroeder, E.B., Shah, S.H., Thacker, E.L., Vanwagner, L.B., Virani, S.S., Voecks, J.H., Wang, N.Y., Yaffe, K., Martin, S.S., 2022. Heart Disease and Stroke Statistics—2022 Update: A Report From the American Heart Association. Circulation 145, E153–E639. https://doi.org/10.1161/CIR.0000000000001052

van de Ven, V., Lee, C., Lifanov, J., Kochs, S., Jansma, H., De Weerd, P., 2020. Hippocampal-striatal functional connectivity supports processing of temporal expectations from associative memory. Hippocampus 30, 926–937. https://doi.org/10.1002/HIPO.23205

van Schie, P.E.M., Becher, J.G., Dallmeijer, A.J., Barkhof, F., Weissenbruch, M.M., Vermeulen, R.J., 2007. Motor outcome at the age of one after perinatal hypoxic-ischemic encephalopathy. Neuropediatrics 38, 71–7. https://doi.org/10.1055/s-2007-984449

Vargha-Khadem, F., Gadian, D.G., Watkins, K.E., Connelly, A., Van Paesschen, W., Mishkin, M., 1997. Differential Effects of Early Hippocampal Pathology on Episodic and Semantic Memory. Science (80-.). 277, 376–380. https://doi.org/10.1126/science.277.5324.376

Waldschmidt, J.G., Ashby, F.G., 2011. Cortical and striatal contributions to automaticity in information-integration categorization. Neuroimage 56, 1791–1802. https://doi.org/10.1016/j.neuroimage.2011.02.011

Wechsler, D., 2009. The Wechsler memory scale-fourth edition (WMS-IV).

Wechsler, D., 1999. Wechsler Abbreviated Scale of Intelligence.

Wilson, B., Ivani-Chalian, C., Aldrich, F., 1991. Rivermead Behavioural Memory Test for Children. Thames Valley Test Co., Bury St. Edmunds, U.K.

Woolley, D.G., Mantini, D., Coxon, J.P., D’Hooge, R., Swinnen, S.P., Wenderoth, N., 2015. Virtual water maze learning in human increases functional connectivity between posterior hippocampus and dorsal caudate. Hum. Brain Mapp. 36, 1265–1277. https://doi.org/10.1002/HBM.22700

Young, J.M., Powell, T.L., Morgan, B.R., Card, D., Lee, W., Smith, M. Lou, Sled, J.G., Taylor, M.J., 2015. Deep grey matter growth predicts neurodevelopmental outcoes in very preterm children. Neuroimage 111, 360–8. https://doi.org/10.1016/j.neuroimage.2015.02.030

