## Supplementary Materials for "Cognitive outcome and its neural correlates after cardiorespiratory arrest in childhood"

### Supplementary Material

**Figure 1. Inclusion and exclusion of patients from the cardiac arrest database.**

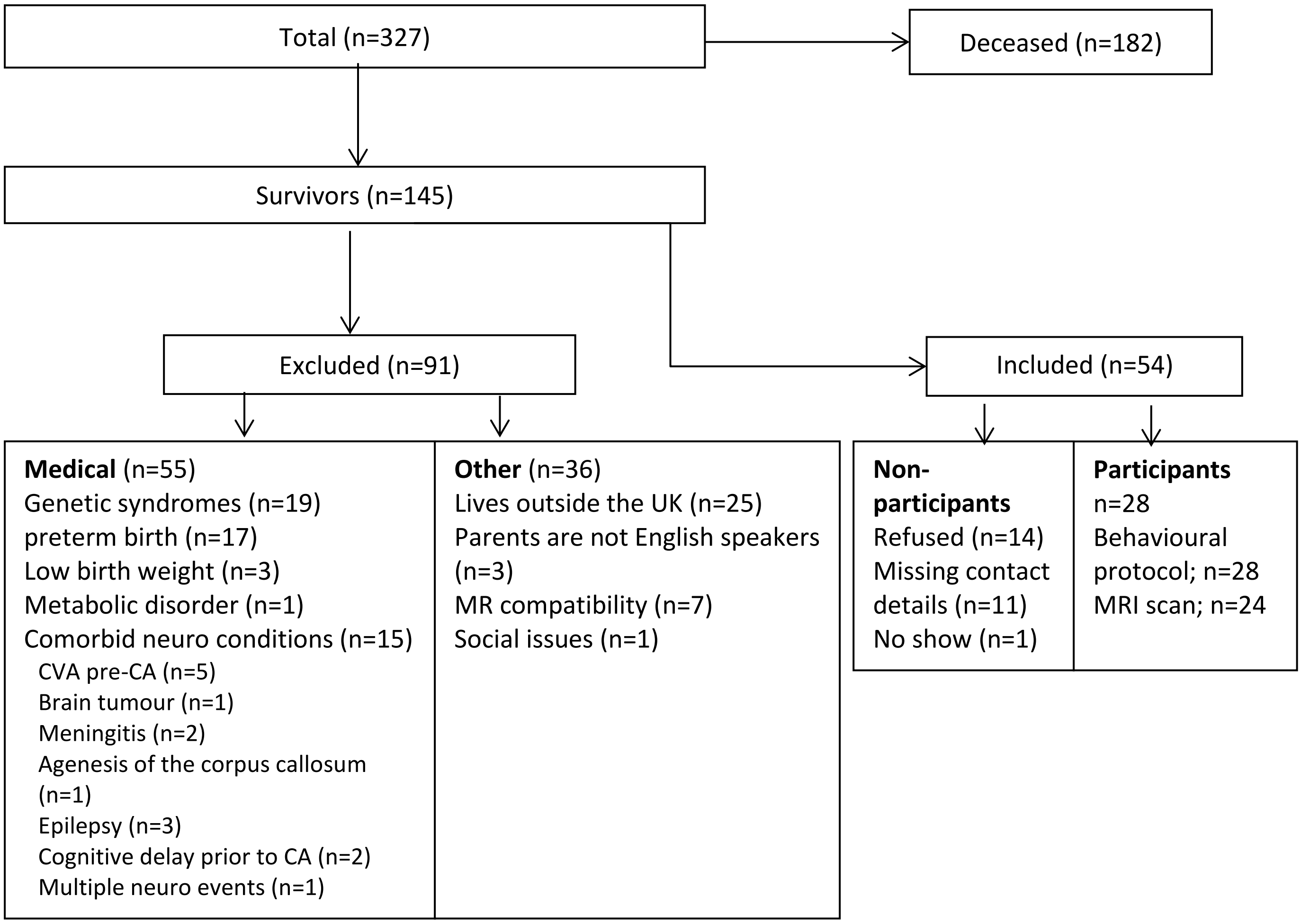

**Table 1. Demographic and clinical information of individual participating patients.**

| ID | Age at test (years) | Gender | Primary aetiology | Diagnosis | CA age (years) | CA time (min) | ECMO duration (days) |
|---|---|---|---|---|---|---|---|
| P01 | 12.3 | M | Respiratory failure | Interstitial lung disease, PH | 3.88 | 7 | 0 |
| P02 | 9.0 | M | Congenital Heart Disease | HLHS | 0.02 | 2 | 0 |
| P03 | 18.0 | F | Sepsis | Septic arthritis ARDS | 13.78 | 1 | 8 |
| P04 | 14.2 | F | Congenital Heart Disease | AVSD | 4.33 | 5 | 0 |
| P05 | 9.4 | F | Congenital Heart Disease | TGA | 0.02 | 3 | 5 |
| P06 | 14.3 | M | Congenital Heart Disease | CoA, VSD | 0.02 | 7 | 0 |
| P07 | 11.2 | F | Congenital Heart Disease | ASD, PH | 10.37 | 7 | 0 |
| P08 | 15.2 | F | Acquired heart disease | Kawasaki disease End-stage heart failure | 13.49 | 23* | 4 |
| P09 | 8.3 | F | Cardiomyopathy | DCM, End-stage heart failure | 0.73 | 4 | 10 |
| P10 | 10.7 | F | Arrhythmia | SVT | 0.05 | 3 | 0 |
| P11 | 10.5 | M | Congenital Heart Disease | CoA | 0.03 | 21* | 0 |
| P12 | 12.3 | F | Cardiomyopathy | DCM, End-stage heart failure | 13.38 | 100 | 10 |
| P13 | 9.0 | M | Respiratory failure | ARDS | 4.92 | 2 | 11 |
| P14 | 18.0 | F | Congenital Heart Disease | VSD, ASD, CoA | 1.44 | 3 | 7 |
| P15 | 14.2 | F | Sepsis | Sepsis | 6.36 | 2 | 0 |
| P16 | 9.4 | M | Cardiomyopathy | DCM, End-stage heart failure | 1.79 | 6 | 9 |
| P17 | 14.3 | M | Congenital Heart Disease | PA, VSD, PDA | 4.07 | 23 | 0 |
| P18 | 11.2 | M | Congenital Heart Disease | TAPVD | 0.04 | 12 | 6 |
| P19 | 15.2 | F | Congenital Heart Disease | Left atrial isomerism, double outlet right ventricle, congenital heart block, End-stage heart failure | 6.79 | 4 | 0 |
| P20 | 8.3 | M | Primary arrhythmia | Primary atrial multifocal arrhythmia, ASD | 0.22 | 7 | 19 |
| P21 | 10.7 | M | Congenital Heart Disease | TGA, VSD, pulmonary stenosis | 2.19 | 43 | 6 |
| P23 | 10.5 | F | Congenital Heart Disease | HLHS | 3.56 | 3 | 0 |

| ID | Age at test (years) | Gender | Primary aetiology | Diagnosis | CA age (years) | CA time (min) | ECMO duration (days) |
|---|---|---|---|---|---|---|---|
| P24 | 15.8 | M | Primary arrhythmia | Wolff Parkinson white syndrome | 0.04 | 4 | 0 |
| P25 | 8.3 | F | Congenital Heart Disease | TGA | 0.02 | 0.5 | 0 |
| P26 | 13.2 | F | Respiratory failure | Fetomaternal Transfusion | 0.00 | 1 | 0 |
| P27 | 13.3 | F | Respiratory failure | Pneumococcal bacterial infection ARDS | 0.82 | 3 | 12 |
| P28 | 14.4 | F | Respiratory failure | Pneumococcal bacterial infection ARDS | 2.92 | 4 | 14 |
| P29 | 12.5 | F | Respiratory failure | Meconium aspiration at birth | 0.00 | 4 | 5 |

* First CA occurred outside hospital. Abbreviations: ARDS = Acute respiratory distress syndrome; ASD = Atrial septal defect; AVSD = Atrioventricular septal defect; CA = cardiorespiratory arrest; CoA = Coarctation of the Aorta; DCM = Dilated Cardiomyopathy; ECMO = Extracorporeal membrane oxygenation machine; HLHS = Hypoplastic left heart syndrome; PA = Pulmonary Atresia; PDA = Patent Ductus Arteriosus; PH = Pulmonary hypertension; TAPVD = Total Anomalous Pulmonary Venous Drainage; TGA = Transposition of the great arteries; VSD = Ventricular Septal Defect.

**Table 2. Comparison of participating and non-participating patients**

| Variable | | Non-participants | Participating Patients | Statistical test | p-value |
|---|---|---|---|---|---|
| **Sex** | Female | 14 | 17 | Chi square | 0.610 |
| | Male | 12 | 11 | | |
| **Age at test (years)**[1] | Mean | 11.46 | 12.04 | t-test | 0.550 |
| | Std. Deviation | 3.67 | 3.34 | | |
| | Range | 7 – 21 | 8 – 20 | | |
| **SES** | Mean | 14,614 | 17,241 | Mann-Whitney | 0.346 |
| | Std. Deviation | 10,700 | 9,367 | | |
| **CA Location** | In Hospital | 26 | 27 | Chi square | 0.331 |
| | Outside Hospital | 0 | 1 | | |
| **Mechanical circulatory support** | No | 25 | 27 | Chi square | 0.957 |
| | Yes | 1 | 1 | | |
| **Heart Transplant** | No | 21 | 22 | Chi square | 0.841 |
| | Yes | 5 | 6 | | |
| **Number of CA** | 1 | 15 | 21 | Chi square | 0.282 |
| | 2 | 7 | 3 | | |
| | 3 | 4 | 4 | | |
| **Age at first CA (days)** | Median | 333.5 | 589.0 | Mann-Whitney | 0.703 |
| | IQR | 1344 | 1729 | | |
| **CA Total Time (min)** | Median | 5 | 4 | Mann-Whitney | 0.531 |
| | IQR | 11 | 4 | | |
| **Days on ECMO** | Median | 0 | 2 | Mann-Whitney | 0.269 |
| | IQR | 5 | 9 | | |

[1] For the non-participant group, age indicates the patient's age on the day when we initially attempted to contact the caregivers for the purpose of study recruitment. Abbreviations: CA = cardiac arrest, ECMO = extracorporeal membrane oxygenation, IQR = interquartile range, SES = socio-economic status.

**Table 3. Radiological findings on MRI.**

| ID | Hippo | Fornix | MB | S > G | BG | Peri-ventricular | Lateral ventricle | Other |
|---|---|---|---|---|---|---|---|---|
| P02 | N | N | N | N | N | N | N | None |
| P03 | N | N | N | N | N | N | N | Old haematoma in genu of CC, R anterior limb of internal capsule and L parietal WM; possible gliotic scar in L deep frontal WM |
| P04 | N | N | N | N | N | N | N | None |
| P05 | N | N | N | N | N | N | N | Slightly small CC |
| P06 | Sm | Sm | Sm | N | N | Slightly reduced, Bilateral gliotic scars | Slightly large | None |
| P07 | N | N | N | N | N | N | N | Possible pituitary haemorrhage; possibly slightly small CC |
| P08 | N | N | N | N | N | N | N | None |
| P10 | N | N | N | N | N | N | N | None |
| P12 | N | N | N | N | N | N | N | None |
| P13 | N | N | N | N | N | N | N | None |
| P14 | N | N | Sm | N | R Caudate injury | Reduced | Dilated | R frontal cortical scar and deep WM gliotic scars |
| P15 | N | N | N | N | N | N | N | None; dental artefacts |
| P16 | N | N | N | N | N | N | N | None |
| P17 | N | N | N | N | N | N | N | None |
| P18 | N | N | N | N | N | N | N | None |
| P19 | N | N | N | S > G | R Caudate & Putamen gliotic scar | N | N | None |
| P20 | N | N | N | N | N | N | N | Small gliotic scar in right frontal WM |
| P21 | N | N | N | N | N | N | N | Bilateral parieto-occipital cortical injury and small optic chiasm and anterior visual pathways |
| P23 | N | N | N | N | N | N | N | None |
| P24 | N | N | N | N | N | N | N | None |
| P25 | N | N | N | N | N | N | N | None |
| P26 | N | Sm | Sm | N | N | N | N | R frontal cortical injury; small CC |
| P27 | N | N | N | N | N | N | N | None |
| P28 | N | N | N | N | L Caudate & Putamen gliotic scar | N | N | R frontal operculum and insula cortex injury; including post central gyrus |
| P29 | N | N | N | N | N | N | N | Slightly small CC |

P01, P09, P11 had no MRI scan. The mesial temporal lobe, dorsomedial thalamus and cerebellum were rated as normal in all patients and are therefore not listed in the Table. Abbreviations: BG = Basal Ganglia; CC = corpus callosum; Hippo = hippocampus; L = left; MB = mammillary bodies; N = normal; R = right; S > G = Splenium > Genu; Sm = small; WM = white matter.

### Table 4. Behavioural scores and Results of Principal Component analyses

#### A. Performance scores on the tests for the patient and the control groups.

| | Control group | | | Patient group | | |
|---|---|---|---|---|---|---|
| | **Mean** | **Standard Deviation** | **Impaired performance** | **Mean** | **Standard Deviation** | **Impaired performance** |
| Memory | | | | | | |
| Sunderland (Z score) | -0.55 | 0.45 | 0 | 0.81 | 1.55 | 0 |
| Verbal Delayed | 113.86 | 10.63 | 0 | 88.52 | 17.81 | 6 (22%) |
| RBMT (% correct) | 92.56 | 6.94 | 0 | 83.05 | 13.04 | 5 (19%) |
| ERRNI Forgetting | 101.74 | 11.48 | 1 (4%) | 97.11 | 12.14 | 2 (7%) |
| **Memory (PCA)** | **0.65** | **0.37** | **0** | **-0.68** | **1.00** | **19 (79%)** |
| Language (ERRNI) | | | | | | |
| Ideas 1 | 112.41 | 12.73 | 0 | 106.93 | 16.98 | 1 (3.5%) |
| Ideas 2 | 115.07 | 12.08 | 0 | 104.71 | 17.83 | 2 (7%) |
| MLU | 110.41 | 15.95 | 1 (4%) | 100.79 | 12.13 | 1 (3.5%) |
| Comprehension | 108.19 | 9.02 | 0 | 98.21 | 14.25 | 4 (14%) |
| **Language (PCA)** | **0.38** | **0.63** | **2 (8%)** | **-0.37** | **1.15** | **10 (36%)** |
| Attainment (WIAT) | | | | | | |
| Numerical Operations | 116.36 | 17.79 | 0 | 89.00 | 23.99 | 9 (33%) |
| Word Reading | 110.16 | 11.79 | 1 (4%) | 94.07 | 18.95 | 6 (22%) |
| Spelling | 107.20 | 13.71 | 1 (4%) | 93.04 | 15.84 | 6 (23%) |
| Reading Comprehension | 116.17 | 8.38 | 0 | 99.27 | 14.01 | 2 (8%) |
| **Attainment (PCA)** | **0.54** | **0.67** | **2 (7%)** | **-0.51** | **1.00** | **13 (50%)** |

Impaired performance [number (percentage) of participants] is defined as follows: Sunderland, PCA components: a score which is more than 1.5 standard deviations below the mean; WIAT, ERRNI, CMS/WMS: scores below 80, which is low or exceptionally low score according to clinical classification; RBMT: impaired range according to manual. Some tests were not completed by all participants.

#### B. Results of Principal Component Analyses

| **PCA 1: Memory** | | **PCA 2: Language** | | **PCA 3: Attainment** | |
|---|---|---|---|---|---|
| *Variance* | 54.41% | *Variance* | 59.9% | *Variance* | 82.1% |
| *KMO* | 0.7 | *KMO* | 0.66 | *KMO* | 0.7 |
| *Bartlett's* $\chi^2$ | 41.5 | *Bartlett's* $\chi^2$ | 83.9 | *Bartlett's* $\chi^2$ | 95.8 |
| *Bartlett's p* | < 0.001 | *Bartlett's p* | < 0.001 | *Bartlett's p* | < 0.001 |
| Sunderland | -0.81 | Ideas 1 | 0.87 | Numerical operations | 0.85 |
| Verbal delayed | 0.85 | Ideas 2 | 0.91 | Word reading | 0.93 |
| RBMT | 0.81 | MLU | 0.51 | Spelling | 0.93 |
| ERRNI Forgetting | 0.39 | Comprehension | 0.74 | | |

ERRNI = The Expression, Reception and Recall of Narrative Instrument; KMO = Kaiser–Meyer–Olkin measure of adequacy; MLU = mean length of utterance; RBMT = Rivermead Behavioural Memory Test; WIAT = Wechsler Individual Achievement Test.
